# Epitope Engineered Human Haematopoietic Stem Cells are Shielded from CD123-targeted Immunotherapy

**DOI:** 10.1101/2023.07.11.547391

**Authors:** Romina Marone, Emmanuelle Landmann, Anna Devaux, Rosalba Lepore, Denis Seyres, Jessica Zuin, Thomas Burgold, Corinne Engdahl, Giuseppina Capoferri, Alessandro Dell’Aglio, Clément Larrue, Federico Simonetta, Julia Rositzka, Manuel Rhiel, Geoffroy Andrieux, Danielle Gallagher, Markus Schröder, Amélie Wiederkehr, Alessandro Sinopoli, Valentin Do Sacramento, Anna Haydn, Laura Garcia-Prat, Christopher Divsalar, Anna Camus, Liwen Xu, Lorenza Bordoli, Torsten Schwede, Matthew Porteus, Jérôme Tamburini Bonnefoy, Jacob E. Corn, Toni Cathomen, Tatjana I. Cornu, Stefanie Urlinger, Lukas T. Jeker

## Abstract

Targeted eradication of transformed or otherwise dysregulated cells using monoclonal antibodies (mAb), antibody-drug conjugates (ADC), T cell engagers (TCE) or chimeric antigen receptor (CAR) cells is very effective for haematologic diseases. Unlike the breakthrough progress achieved for B cell malignancies, there is a pressing need to find suitable antigens for immunotherapy of myeloid malignancies. CD123, the interleukin-3 (IL-3) receptor alpha-chain, is highly expressed in various haematological malignancies, including acute myeloid leukaemia (AML) and blastic plasmacytoid dendritic cell neoplasm (BPDCN). However, shared expression of CD123 on healthy haematopoietic stem and progenitor cells (HSPCs) bears the risk for extensive myelotoxicity upon targeted depletion. Here, we demonstrate that rationally designed, epitope-engineered HSPCs were completely shielded from CD123-targeted immunotherapy but remained fully functional while CD123-deficient HSPCs displayed a competitive disadvantage. Thus, molecularly shielded HSPCs could allow tumor-selective targeted immunotherapy and in parallel enable rebuilding a fully functional haematopoietic system. We envision that this approach is broadly applicable to many targets and cells, could render hitherto undruggable targets accessible to immunotherapy and will allow continued posttransplant immunotherapy, for instance to treat minimal residual disease (MRD) or be used as a salvage therapy. Since the function of the engineered targets is preserved, multiplexed molecular shielding could also enable targeted combination immunotherapies to address tumor heterogeneity. More generally, epitope shielding will be applicable for replacement of other cell types including the many immune cells which are currently being considered for engineered cellular therapies.

## Introduction

Targeted cell depletion represents a medical standard of care for several liquid malignancies, autoimmune diseases and prevention or treatment of acute rejection in organ transplantation. Depleting antibodies are mostly IgG but other highly effective targeted immunotherapies work through different mode of action (MoA) and include various antibody-derived molecular formats such as antibody-drug conjugates (ADC), radioimmunoconjugates, T cell engagers (TCE) or chimeric antigen receptor (CAR) bearing cells^1–3^. In the past decade the latter have emerged as a highly effective, programmable cell depletion modality with very high response rates in B cell malignancies and more recently systemic lupus erythematodes^4–9^. The persistence of CD19 CAR T cells can be reliably measured by assessing the duration of B cell aplasia after infusion ^5,10,11^. In children and young adults with B-ALL it appears that persistence of CAR T cells is an important requirement for cure^12^. Hence, various strategies to increase CAR T longevity are explored^13,14^. Although high depletion efficiencies can lead to long-term remission, they are associated with the risk for prolonged B cell aplasia due to indiscriminate killing of both normal and tumor B cells, respectively^9,15^. In fact, CAR T cells can persist for a decade and result in years-long B cell aplasia^16^. Such deep purging of a cell type is only acceptable if the depleted cell is dispensable and/or its function can be replaced. Thus, a key to the CAR T field’s success was the availability of B cell-restricted target antigens (e.g. CD19) and the possibility to mitigate the loss of co-targeted healthy B cells through infusions of immunoglobulins (IVIG). Given the clinical benefit and the commercial availability of CAR T cells as well as other effective cell depleting modalities, the number of patients at risk for iatrogenic long-term immunodeficiencies through highly efficacious cell depleting therapies are expected to continue to increase rapidly.

Unlike the breakthrough progress achieved for B cell malignancies there is a pressing need to find suitable antigens for immunotherapy of myeloid malignancies and particularly acute myeloid leukaemia (AML)^17^. Targeting myeloid malignancies is especially challenging however, since AML displays high clonal heterogeneity, being composed of cells with highly variable surface protein expression^12,17–19^. Importantly, leukaemia stem cells (LSCs) which are phenotypically very similar to HSCs are important targets in (AML)^20^. Therefore, most AML candidate targets are co-expressed by HSPCs^17,18^. As a consequence, the risk of myelosuppression associated with myeloid-cell targeted CAR T therapy likely limited the number of clinical trials for AML compared to B cell targeted CAR T trials^12,21^. Despite major efforts, single targets with an expression profile as favourable as CD19 could not be identified in AML^17^. Therefore, the paucity or mere absence of cancer-restricted surface proteins constitutes a critical barrier to antigen-specific immunotherapy^11,12,22^.

A number of solutions have been proposed to overcome the limitations of shared target antigens that underly on-target off-tumour toxicity. Affinity tuning, i.e. reducing a CAR’s affinity, can increase the selectivity towards cells with a high antigen expression^23^; transient CAR expression (delivery as mRNA) temporally limits CAR activity; and CAR T therapies targeting essential cells, particularly HSPCs, could be used as a bridge to transplant^21,24,25^. However, sparing cells with low antigen expression creates a risk for antigen^-/lo^ relapse^12,26^ and the need to remove an effective CAR T therapy targeting HSPCs is neither economically nor scientifically appealing. Therefore, three groups proposed a radical solution: removing the target antigen partially or entirely in HSPCs before haematopoietic stem cell transplantation (HSCT) prevents binding of the immunotherapy to the HSPCs and their progeny and thus creates a synthetic tumor-selectivity^27–29^. Preclinical studies demonstrate the feasibility for CD33 and early results from a clinical trial showed a favourable safety profile with engraftment of CD33-deficient CD34^+^ HSPCs (clinicaltrial.gov NCT04849910)^27–29^. Additional data will be required for further development of this approach. However, the number of truly dispensable antigens is likely limited yet mostly unknown. Moreover, targeting dispensable proteins may favour antigen negative cancer relapses, a phenomenon known to limit long-term outcome of CAR T therapies^26^. Therefore, deliberately targeting essential proteins would be preferrable to reduce the risk for antigen escape. However, this is impossible with a knock-out (KO) approach.

Here, we aimed to provide proof-of-concept by engineering HSPCs expressing an endogenous protein variant that completely shields the cells from targeted immunotherapy while preserving its function. The interleukin-3 receptor α chain (IL3RA; CD123) regulates proliferation and differentiation of HSPCs^30^, is often expressed on AML LSCs and blasts from relapses^31^ and hence is associated with a poor outcome of the disease. Therefore, CD123 constitutes a promising target for AML^19,21^ for which multiple therapeutics are being explored preclinically and clinically including IL-3 bound to diphtheria toxin^32^, monoclonal antibodies (mAb) blocking IL-3 ^33^ or engineered for enhanced antibody dependent cellular cytotoxicity (ADCC)^34^, an antibody drug conjugate (ADC)^35^ and bispecific TCEs, e.g. a CSL362/OKT3-TCE^36,37^. Despite the promising on target toxicity, many of these therapies displayed cytotoxicity towards HSPCs, monocytes, basophils and plasmacytoid dendritic cells (pDCs)^19,21,36^. Thus, protecting HSPCs from the immunotherapies is desirable but due to CD123’s role in HSPC biology it is unknown whether a CD123 KO approach would be viable or if a KO approach may result in impaired immune function. We identified multiple single amino acid (aa) substitutions that shielded from ADCC, ADC, TCE and CAR T cell killing while preserving CD123 function. These variants enabled selective TCE- or CAR T-mediated tumour killing while engineered HSPCs were unaffected. CD123 KO HSPCs had a major competitive disadvantage in vitro while engineered CD123 knock-in cells were comparable to wt. Given the favourable safety data, we envision that this approach is broadly applicable, could render undruggable targets accessible to immunotherapy and will allow continued, posttransplant immunotherapy for instance to treat minimal residual disease (MRD) or be used as a salvage therapy.

## Results

CD123 variants were designed *in silico*. We aimed to identify protein variants that are structurally and functionally tolerated, i.e., variants that exhibit a similar structure to wildtype (WT) CD123, that preserve the ability to bind IL-3 and elicit IL-3 mediated downstream cell signalling but are otherwise shielded from the CSL362 antibody or a similar antigen binding moiety. To this aim, available experimentally determined three-dimensional structures of CD123 in different states were used to identify the protein regions involved in binding to CSL362 and IL-3. As previously described^38^, the CSL362 epitope is located at the N-terminal domain (NTD) of CD123 (**Fig. 1a**), where the CSL362 fragment antigen binding (Fab) binds both the “open” and “closed” conformations of the NTD. Based on structural analysis, we identified several CD123 residues as part of the antibody-antigen interface and involved in direct intermolecular interactions with the CSL362 complementarity-determining regions (CDRs) in both conformational states: T48, D49, E51, A56, D57, Y58, S59, M60, P61, R84, V85, A86, N87, P89, F90, S91. Most of these amino acid sites show differential exposure to solvent upon binding of the CSL362 antibody, with residues E51, S59, P61, and R84 switching from highly exposed to buried (**Fig. 1b**). E51, S59 and R84 were selected as critical sites for the CSL362 binding (**Fig. 1c**) and most likely safe for mutagenesis. In contrast, P61 is likely relevant for IL-3 binding due to its close interatomic distance to IL-3 residues and therefore was excluded from further analysis. Comprehensive mutagenesis was performed *in silico* and the mutational effect estimated as statistical energy difference (ΔE) (**Fig. 1d**) with particular focus on the three key residues (**Fig. 1e**) and candidate variants selected upon ranking for decreasing ΔE, for a total of 28 variants (**Fig. 1f**).

**Figure 1.**
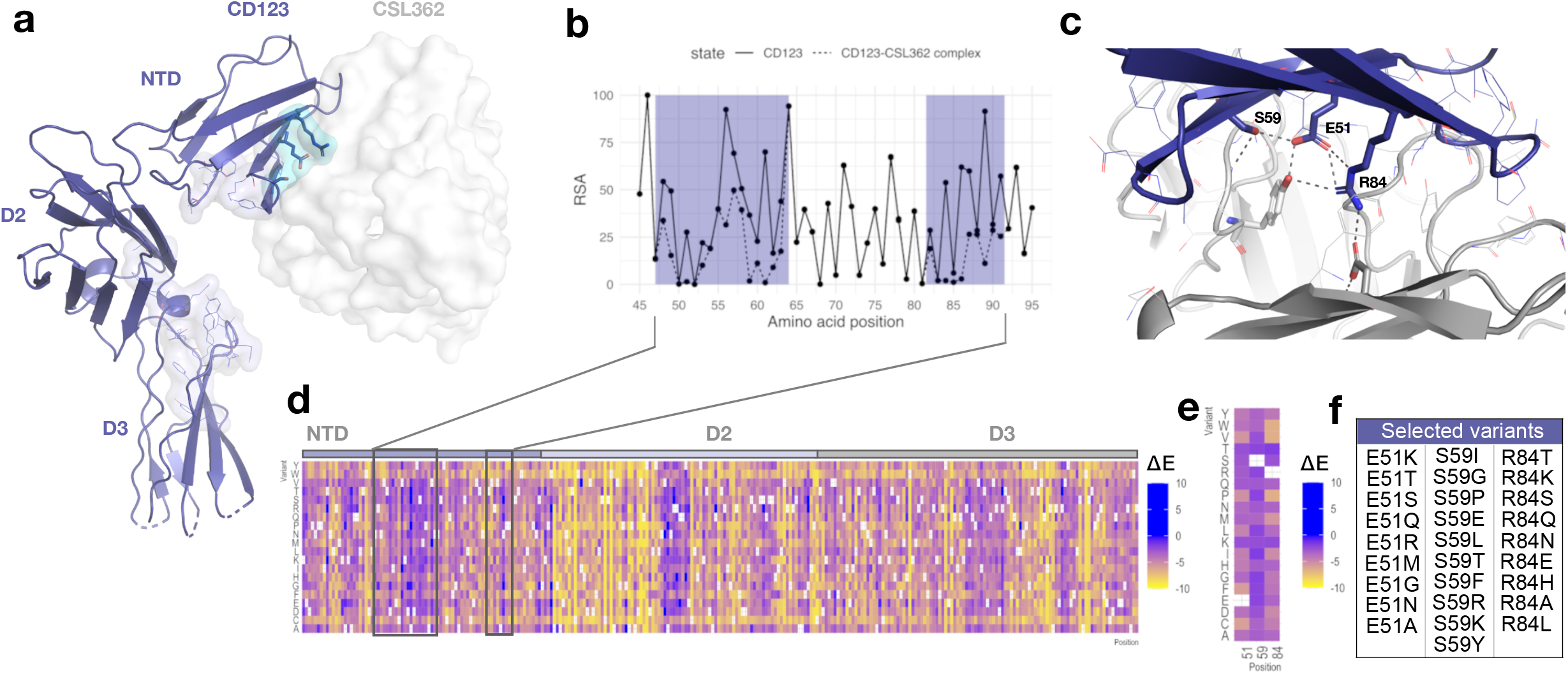
Rational design of human CD123 protein variants to shield from targeted immunotherapy. **a**, Crystal structure of the CD123-CSL362 complex in open conformation (PDB ID: 4JZJ^38^). CD123 is shown as ribbons. The CSL362 antibody variable domain is shown as white surface. CD123 amino acid residues involved in IL-3 binding are highlighted as lines and in light-blue. **b**, Per-residue relative solvent accessibility (RSA) computed on the CSL362-free (solid line) and CSL362-bound (dashed line) states based on the X-ray structure of the CD123-CSL362 complex (PDB ID: 4JZJ). RSA data are shown for the N-terminal domain. The CSL362 epitope region is highlighted by blue rectangles. **c**, Amino acid residues at the interface of the CD123-CSL362 complex are highlighted as lines and sticks. Side chain mediated intermolecular contacts are shown as dashed black lines. **d**, The predicted ΔE mutational landscape of CD123 is shown as a heatmap for the full extracellular domain (residue range 20-305, x-axis) and **e**, selected amino acid positions: E51, S59 and R84. Heatmap color ranges from yellow (ΔE <0, predicted damaging) to blue (ΔE ≥0, predicted neutral or beneficial). **f**, Selected amino acid variants at residues E51, S59, R84 sorted by decreasing ΔE values.

We used a CSL362 IgG1 biosimilar (MIRG123) to experimentally validate its binding to the in silico designed CD123 variants. We generated HEK-293 cells stably expressing human wildtype CD123 (HEK-CD123) or individually harbouring each of the 28 selected amino acid substitutions at positions E51, S59 and R84 (**Fig. 1f**). Using flow cytometry we concomitantly quantified MIRG123 binding as well as preserved expression of the variants by staining with the control anti-CD123 mAb clone 6H6, whose binding to CD123 does not interfere with MIRG123 (**Fig. 2a**). The results revealed a drastic reduction of MIRG123 binding to most candidate CD123 variants, and abolished binding to almost half (13/28) of them. Based on the dual staining characteristics to mAbs 6H6 and MIRG123 each variant was categorized as either a non-binding (< 1% double staining to 6H6/MIRG123, 13 variants, blue), weak (1-20%, 12 variants, orange) or strong (> 20%, 3 variants, red) binding variant (**Fig. 2b**). The latter showed comparable binding to HEK-CD123. Among the weak binding variants, MIRG123 mostly bound the highest CD123 expressing cells. These results demonstrate the feasibility to rationally design single amino acid substitutions that completely disrupt the binding of a candidate antibody while preserving the expression of the engineered protein.

**Figure 2:**
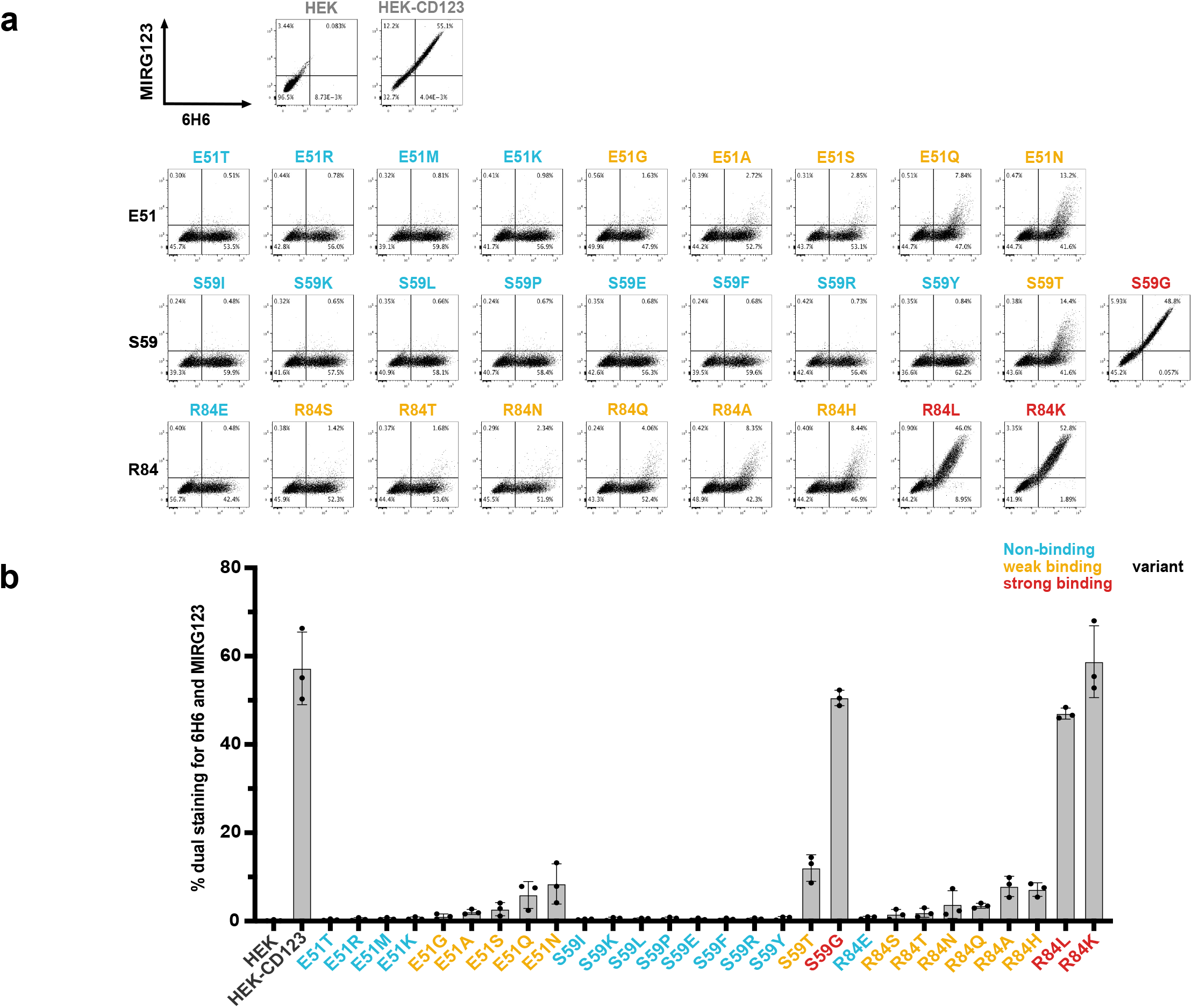
Preserved expression of engineered CD123 variants despite abolished binding to the monoclonal antibody MIRG123. **a,** Flow cytometry plots and **b,** summarizing bar graph showing binding of the anti-human CD123 antibody MIRG123 (biosimilar of CSL362) and the control clone 6H6 to wildtype CD123 and its 28 variants stably expressed in HEK-293 cells. Variants were categorized based on the dual staining to MIRG123 and 6H6 as non-binding (blue, <1% dual staining), weak (orange, 1-20%) or strong (red, >20%) binding variants. Control conditions (grey) are HEK-293 cells stably expressing wildtype CD123 (HEK-CD123) and non-transduced HEK-293 cells (HEK). Error bars: mean ± SD. Each symbol represents an individual experiment. Data in **a** is representative of three independent experiments.

Next, we explored whether the non-binding variants were protected from MIRG123-mediated cytotoxicity either in the format of a mAb (ADCC), a bispecific TCE or human CAR T cells (**Fig. 3a**). First, we tested whether the CD123 variants were shielded from MIRG123-triggered ADCC using an FcγRIIIa expressing reporter cell line. Target cell binding of the test antibody leads to activation of a luminescence signal in the reporter cells through Fc-mediated activation of the FcγRIIIa receptor. High luminescence was measured in the presence of unmodified antigen (HEK-CD123) as well as all strong binding variants (**Fig. 3b**). In contrast and consistent with the flow cytometry data (**Fig. 2a-b**) 12/13 non-binding variants did not induce any ADCC signal above background detected in control cells devoid of CD123 (HEK; lower dashed line) with the exception of S59Y which induced minimal activity. Weak binders showed an intermediate ADCC activity. Thus, ADCC signals positively correlated with the flow cytometry data.

**Figure 3:**
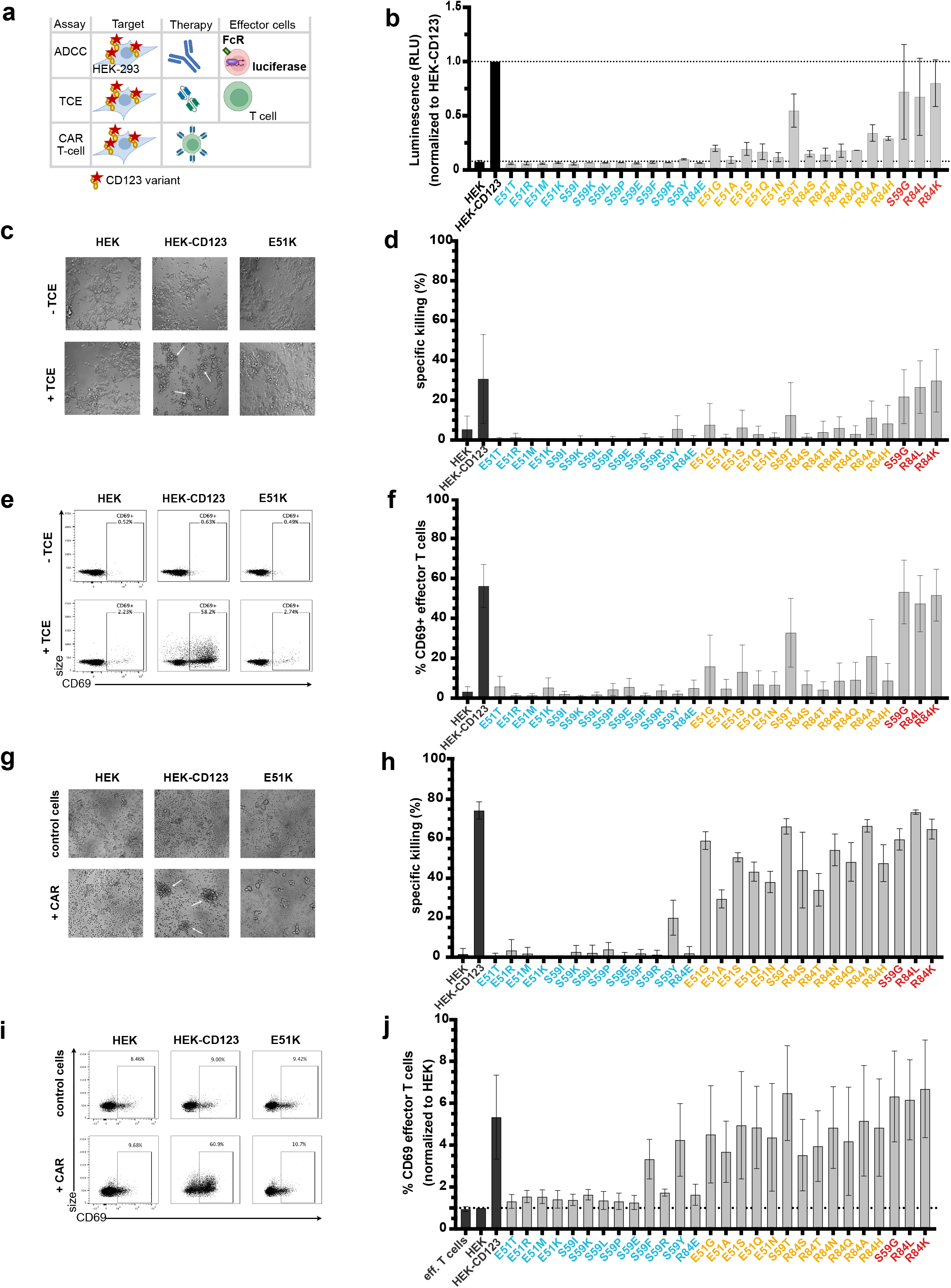
Cells expressing engineered CD123 variants are shielded from multiple targeted immunotherapy modalities *in vitro*. **a,** Schematic to assess shielding of CD123 expressing cells from three targeted immunotherapies (ADCC, TCE, CAR T-cell) in vitro. **b,** MIRG123-induced ADCC measured by luminescence of the effector cell line Jurkat/FcγRIIIa/NFAT-Luc following co-culture with HEK, HEK-CD123 or the CD123 variants. Luminescence signal normalized to the culture with HEK-CD123 (top dashed line). Data of 2 independent experiments. **c-f,** 3-day co-culture of effector T cells and HEK-293 expressing CD123 and its variants with and without CSL362/OKT3-TCE (TCE). Data represent 5 independent blood donors and experiments with 2 technical replicates per group. **c,** Representative images (Magnification 20x) after 3-day co-culture with HEK, HEK-CD123 or E51K with and without TCE. White arrows indicate cell clustering. **d,** Specific TCE-mediated killing of HEK-293 cells or its variants. **e,** Representative flow cytometry plots indicating CD69-expression in effector T cells without (top) and with (bottom) TCE after 3 days co-cultures with HEK, HEK-CD123 and the variant E51K. **f,** Frequency of CD69-expressing CD3^+^ effector T cells after 3 days. **g-j,** Human 123CAR T cells were co-cultured with the target cells HEK, HEK-CD123 or its variants for 24h. Control T cells were electroporated with an HDRT, but incomplete RNP. Data from 3 independent blood donors and experiments with 2 technical replicates per group. **g,** Representative microscopy images (Magnification 20x) after 1 day with white arrows indicating cell clustering. **h,** Specific killing of target cells measured by flow cytometry at day 1 of co-culture. **i,** Representative FACS plots and **j,** summary of CD69^+^ 123CAR T cells either alone (effector T cells) or in the presence of HEK, HEK-CD123 or all CD123 variants after 24h co-culture. The data are normalized to %CD69^+^cells in the presence of HEK target cells. **b-j,** Error bars: mean ± SD.

Encouraged by these results, we investigated whether the non-binding variants would also protect from more potent immunotherapies such as TCEs and CAR T cells which result in cytotoxicity at a lower antigen density than ADCC-inducing mAbs^39^. First, we co-cultured CD123 expressing HEK-293 cells and human T cells with and without the CSL362/OKT3-TCE (TCE)^37^. After 3 days of co-culture, T cells clustered in the presence of HEK-CD123 and TCE, whereas clusters were neither seen with the non-binding variant E51K (or any of the other non-binding variants) nor with the target cells devoid of CD123 (HEK) (**Fig. 3c**). In line with these findings, neither HEK cells nor cell lines expressing non-binding CD123 variants were subject to TCE-mediated cytotoxicity. In contrast, some weak binding CD123 variants induced limited cytotoxicity while control HEK-CD123 and strong-binding variants were lysed in the presence of the TCE (**Fig. 3d**). Consistent with these results, upregulation of the activation marker CD69 above background was only detected in T cells in co-culture with HEK-CD123 but not with E51K or HEK (**Fig. 3e**) or any of the non-binding CD123 variants (**Fig. 3f, Extended data Fig. 1a**). In contrast, the frequency of CD69^+^ T cells increased in the presence of some weak binding variants and was comparable between HEK-CD123 and the strong binding variants. Both CD4^+^ and CD8^+^ T cells were activated but CD8^+^ T cells displayed a higher proportion of activated CD8^+^CD69^+^ effector T cells (**Extended data Fig. 1b-c**). Likewise, secreted IFNγ was only detected in the co-culture supernatants from highly activated T cells triggered by strong antigen binding (**Extended data Fig. 1d**).

As a third modality of CD123-targeting immunotherapies we investigated CAR T cells. We used non-viral CRISPR/Cas9-mediated homology directed repair (HDR) to integrate a second-generation CSL362-derived CD123-specific CAR (123CAR) into the T cell receptor α constant region (*TRAC*) (**Extended data Fig. 2a**). Up to 9% of all electroporated CD4^+^ and CD8^+^ T cells expressed the 123CAR as assessed by GFP (**Extended data Fig. 2b-d**). Correct 123CAR integration at the *TRAC* locus was confirmed by Sanger sequencing (**Extended data Fig. 2e**). GFP^+^123CAR T cells were purified by flow cytometry, expanded, and subsequently used for in vitro cytotoxicity assays. CAR-less control T cells or 123CAR T cells were co-cultured for 24h with the CD123 variant expressing target cell lines. Within hours, 123CAR T cells clustered around HEK-CD123 cells, whereas neither HEK nor E51K triggered 123CAR T clustering (**Fig. 3g**). As observed for ADCC and TCE, 12/13 non-binding CD123 variants were shielded from cytotoxicity (**Fig. 3h**). The only exception was again S59Y which induced cytotoxicity (**Fig. 3h**) and IFNγ secretion (**Extended data Fig. 2i**). In contrast and different to the results observed for ADCC and TCE, both, weak and strong binding variants triggered strong cytotoxicity (**Fig. 3h**) and IFNγ secretion (**Extended data Fig. 2i**). In accordance, neither HEK nor E51K increased the proportion of activated CD69^+^ 123CAR T cells above background whereas the frequency of CD69^+^ 123CAR increased in response to unmodified CD123 (**Fig. 3i**). The exceptions were S59F and S59Y which both induced CD69 upregulation in CD4^+^ and CD8^+^ 123CAR T cells, respectively (**Fig. 3j; Extended data Fig. 2f-h**). However, among the non-binders, only S59Y concomitantly induced CD69 upregulation, IFNγ secretion and cytotoxicity whereas S59F resulted in isolated CD69 upregulation but neither IFNγ secretion nor cytotoxicity. Thus, in contrast to ADCC and TCE where antibody and T cell effector function correlated well with antibody binding as determined by flow cytometry (**Fig. 3b-f**, **Fig. 2b**), weak binding CD123 variants induced 123CAR T responses comparable to strong-binding CD123 variants. These findings suggest that CAR T cells are more potent than ADCC and TCE and even weak residual binding can result in CAR T activation and elimination of target cells. In summary, we demonstrate that a series of single amino acid substitutions in CD123 are sufficient to completely prevent the activity of an ADCC-inducing mAb, a TCE and a CAR T cell. This suggests that such variant proteins - when engineered into the genome of target cells - might provide protection from different targeted immunotherapy.

To characterize the effect of the introduced point mutations on the protein biophysical properties, the extracellular domains (ECD, amino acids 19-305) of selected CD123 variants were synthesized as soluble proteins. Real time interaction with the immobilized CSL362 was measured using Bio-Layer Interferometry (BLI) at increasing concentrations of soluble CD123 ECD variants. Wildtype CD123 associated rapidly to the antibody CSL362 in a dose-dependent manner (**Fig. 4a**). In contrast, no interaction was observed up to 300nM of the soluble analyte CD123 E51T, a variant characterized as a non-binder by flow cytometry (**Fig. 4a**, **Fig. 2a, b**). Similarly, no association to CSL362 was observed for other non-binding variants, including E51K, S59P, S59E, S59F, S59R and R84E. Residual CSL362 association was detected at different concentrations of the weak-binding variants E51A, E51Q, R84T and R84Q (**Fig. 4b**). Consistent with the shielding assays (**Fig. 3b-j**), S59Y showed similar residual binding as weak-binders. Importantly, the association to the control antibody 6H6 was preserved in all tested variants, except for R84E (**Fig. 4c**). Next, we assessed the functionality of the variants in terms of binding to IL-3, i.e. the physiologic ligand of CD123, to the immobilized CD123 variants. All tested variants bound to increasing concentrations of IL-3, with S59P and R84E displaying a modest reduction in IL-3 binding (**Fig. 4d**). Lastly, protein thermal stability was assessed by differential scanning fluorimetry (DSF) analysis. Most CD123 variants demonstrated a thermostability comparable to CD123 WT with an unfolding temperature (Tm) of 50°C. In contrast, R84E induced a higher fluorescence signal at ambient temperature and featured a lower Tm (43°C) (**Fig. 4e**). Since R84E showed reduced binding to 6H6, IL-3 and a lower thermal stability compared to wildtype, it was excluded from further experiments. In summary, we identified several protein variants harbouring single amino acid substitutions at two different residues (E51T, E51K, S59E, S59R) that entirely abolish binding to MIRG123 but preserve protein stability, binding to the control mAb 6H6 and the receptor’s natural ligand IL-3.

**Figure 4.**
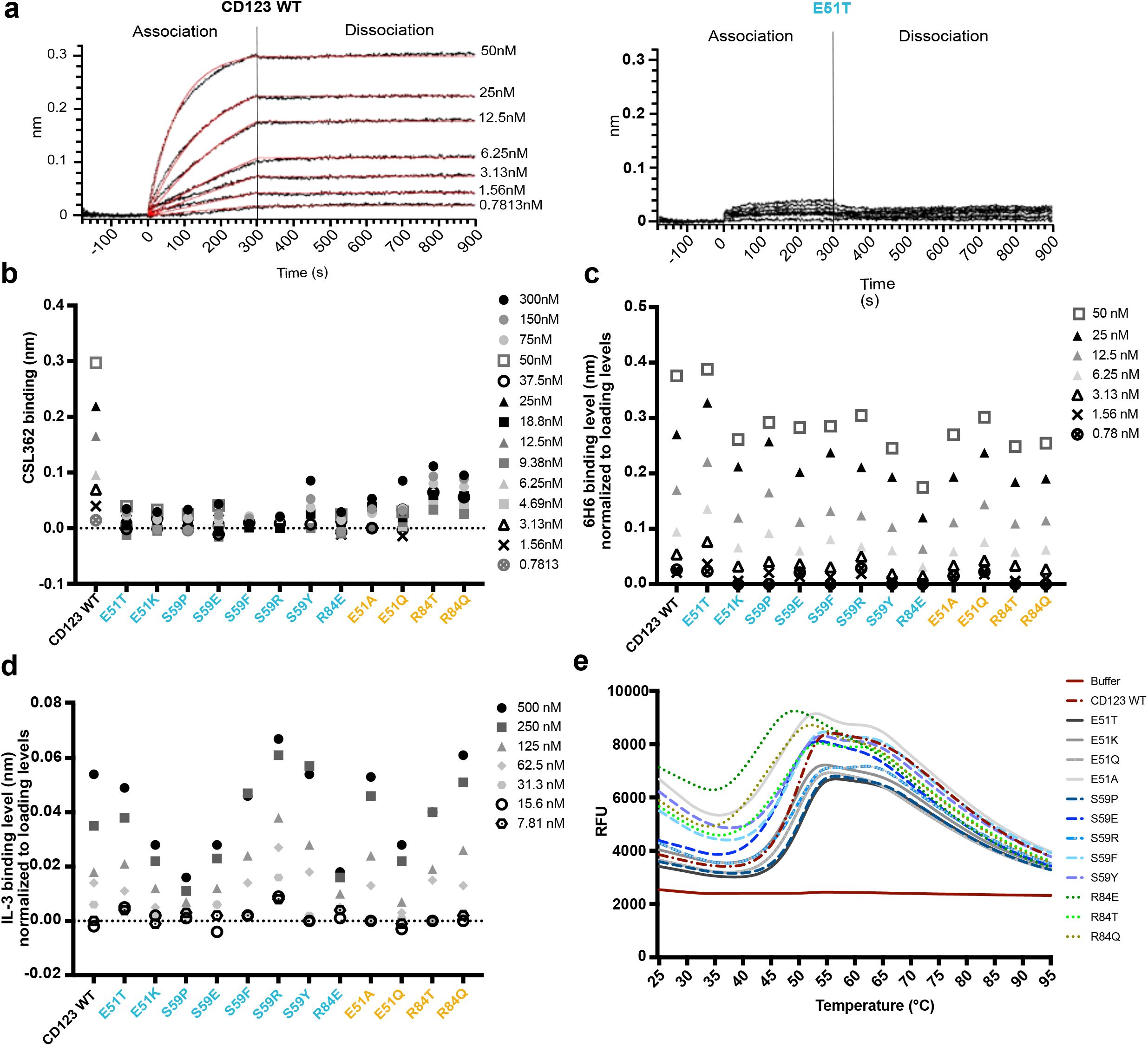
Biophysical characterization of selected CD123 protein variants. **a,** Binding of CSL362 to recombinant extracellular domain (ECD) of CD123 WT (**left**) and CD123 E51T (**right**) at increasing concentrations measured of CSL362 by Bio-Layer Interferometry. **b**, Binding levels of CD123 WT and its variants at different concentrations to captured CSL362 at 280s. CD123 WT reaches its saturation to CSL362 at 50nM, therefore higher concentrations were not measured. **c,** Binding levels of CD123 WT and its variants to the captured antibody 6H6 (normalized to the loading level of 6H6) at 250s. **d,** Binding levels of IL-3 to biotinylated CD123 WT and variants (normalized to loading levels of biotinylated CD123 WT and its variants) at 250s. **e,** Thermal unfolding (Relative Fluorescence Unit, RFU) of CD123 WT and variants measured by Differential Scanning Fluorimetry (DSF) with increasing temperatures.

Since we used a cell-free system to establish that the selected CD123 variants preserved dose-dependent binding of IL-3, we next aimed to investigate the functionality of E51K and E51T in intact cells. We used CRISPR/Cas9-mediated HDR to engineer the variants into the human erythroleukemia cell line TF-1 whose proliferation and survival is IL-3- or GM-CSF-dependent^40^ (**Extended data Fig. 3a**). The genetically engineered cells were cultured with increasing concentrations of IL-3. WT TF-1 displayed IL-3-dependent growth and the two KI populations E51K and E51T demonstrated near overlapping growth curves indicating intact IL-3 sensing and signalling. In contrast, KO sorted cells proliferated considerably less (**Extended data Fig. 3b**). To test the blocking effect of MIRG123, we cultured the cells with a fixed IL-3 concentration but increasing amounts of MIRG123. As reported in a comparable system using BaF3 cells^38^, WT cells showed a dose-dependent growth inhibition and died in the presence of antibody concentrations ≥ 0.036nM. In contrast, E51K and E51T KI cells proliferated irrespective of the MIRG123 concentration. KO cells were not affected by MIRG123 but displayed decreased survival compared to WT, E51K or E51T (**Extended data Fig. 3c**).

Next, we aimed to engineer clinically relevant HSPCs that can potentially be used to provide a patient with a CD123 immunotherapy-resistant haematopoietic system. We used reagents close to GMP-grade with the goal to provide scientific proof of concept with the potential for clinical translation. To this end, we engineered the variants E51K and E51T into mobilized, CD34^+^ enriched peripheral blood HSPCs from healthy donors. CD34^+^ HSPCs were electroporated with high fidelity SpCas9 RNPs and single strand oligodeoxynucleotides (ssODNs). CD123 expression was monitored using flow cytometry with the mAbs MIRG123 and 6H6, two and five days post electroporation, respectively (**Fig. 5a, Extended data Fig. 3d**). Preliminary experiments demonstrated that the ssODNs reduced HSPC viability and consequently in vivo engraftment when E51K and E51T HSPCs were compared to HSPCs that were electroporated without a ssODN. Therefore, we designed ssODNs containing the same silent mutations as in the E51K and E51T ssODNs (wt template) and ssODNs containing stop codons in all 3 reading frames (KO template) in order to have similar ssODNs in the control samples. These groups were compared to E51K and E51T KI HSPCs as well as EP only and RNP without ssODN (KO). In comparison to electroporated WT cells (EP), cells receiving RNPs without ssODN displayed only a slightly increased fraction of MIRG123^-^6H6^-^ (KO) cells (**Fig. 5a**). In contrast, HSPCs receiving RNP together with the wt template displayed increased CD123 KO, an effect that was further enhanced in the KO template group as has been reported before. The E51K or E51T HSPCs displayed a distinctly different FACS profile. Over time a population with retained 6H6 but abolished MIRG123 binding (MIRG123^-^6H6^+^) gradually appeared (average 10% on day 2; average 21% on day 5 across 8 donors), suggesting that these were E51K or E51T KI HSPCs (**Fig. 5a, b**). Importantly, CD34^+^CD38^-^CD90^+^CD45RA^-^ cells (HSCs) displayed similar editing rates as the bulk CD34^+^ HSPCs suggesting that long-term repopulating HSCs (LT-HSC) and multipotent progenitors MPP1 and MPP2 were edited at similar frequencies (**Fig. 5c, Extended data Fig. 3e**). Correct editing of E51 (GAG) to K (AAG) or T (ACC) was confirmed by next generation amplicon sequencing (Amplicon-NGS) (**Fig. 5d**). The number of KI reads correlated well with the FACS analysis (**Fig. 5a, b, d**). To investigate whether the variants E51K and E51T preserved IL-3 signalling we analysed pSTAT5 in KI HSPCs and compared them to HSPCs engineered with wt template or KO template. Among CD123^+^ wildtype cells (MIRG123^+^6H6^+^ “wt”, left panel) IL-3 induced a bi-modal pSTAT5 signal in both control HSPCs (wt template) as well as E51K and E51T KI HSPCs (**Fig. 5e, histograms**). Within MIRG123^-^6H6^-^ CD123 KO cells (KO template) pSTAT5 signalling was abolished (**Fig. 5e, histogram**). In contrast, gating on MIRG123^-^6H6^+^ (KI) cells demonstrated that all E51K and E51T KI cells were pSTAT5^+^. Thus, we leveraged the ability to discern unedited from edited cells by flow cytometry to demonstrate that all phenotypically defined KI cells transmitted the IL-3 signal. To further characterize the functionality of the engineered HSPCs and to determine the functional relevance of a preserved IL-3/CD123 signalling axis, we analysed the HSPC in vitro differentiation potential in a colony forming assay. Compared to HSPCs that were electroporated only (EP), the number of colonies was reduced in all samples electroporated with a ssODN template (**Fig. 5f**). Among the latter, HSPCs electroporated with the KO template formed the lowest number of colonies. In contrast, E51K and E51T HSPCs formed equal (E51K) or slightly increased (E51T) number of colonies when compared to wt template HSPCs. However, the relative distribution of colonies representing different lineages (erythroid (E) and myeloid colonies (M)) was comparable among all genotypes (**Fig. 5f**). These results suggested that CD123-deficient HSPCs could have a competitive disadvantage compared to WT or KI HSPCs. To verify this, we analysed the allele frequency (genotypes) found in individual single cell-derived colonies and categorized the alleles as “wt” (unedited), “KO” (NHEJ), “ki” (identification of knock-in template) (**Fig. 5g**). In colonies derived from “wt template” we found only paired alleles with the ability to express intact CD123, i.e. indels were only found when the other allele was wt or ki, and no KO/KO sequences were identified. In KO template HSPCs only very few colonies harboured a genotype leading to the absence of CD123 (KO/KO, ki/ki or ki/KO), whereas the majority of colonies were unedited or contained at least one wt allele (wt/wt, wt/KO or wt/ki). Similarly, in E51K and E51T HSPCs we did not find any KO/KO colonies. In contrast, ki/ki colonies were found at least with equal frequency when compared to wt/wt. Thus, there was a strong counterselection against CD123-deficient colonies whereas CD123 KI sequences were found at the expected frequencies. Therefore, the defect of CD123-deficient colonies in Fig. 5f is an underestimate and the real competitive disadvantage of CD123 KO HSPCs is much more pronounced. In contrast, CD123 E51K and E51T KI HSPCs are functionally equivalent to wildtype with regards to colony forming and differentiation potential. This was confirmed by a flow cytometry-based in vitro differentiation assay as all tested genotypes displayed an equal differentiation into CD33^+^ myeloid and GlycophorinA (GlyA)^+^ erythroid cells (**Fig. 5h**). Of note, also in the presence of CSL362 conjugated to tesirine via an uncleavable linker (CSL362-ADC) HSPCs differentiated normally in myeloid and erythroid lineages. Finally, we evaluated the resistance of myeloid CD33^+^, CD14^+^ or CD15^+^ cells derived from CD123 engineered HSPCs to CSL362-ADC. CD123 expressing 6H6^+^ cells were strongly depleted in EP, wt and KO samples (**Fig. 5i**), whereas E51K and E51T edited cells were largely resistant to CSL362-ADC killing.

**Figure 5.**
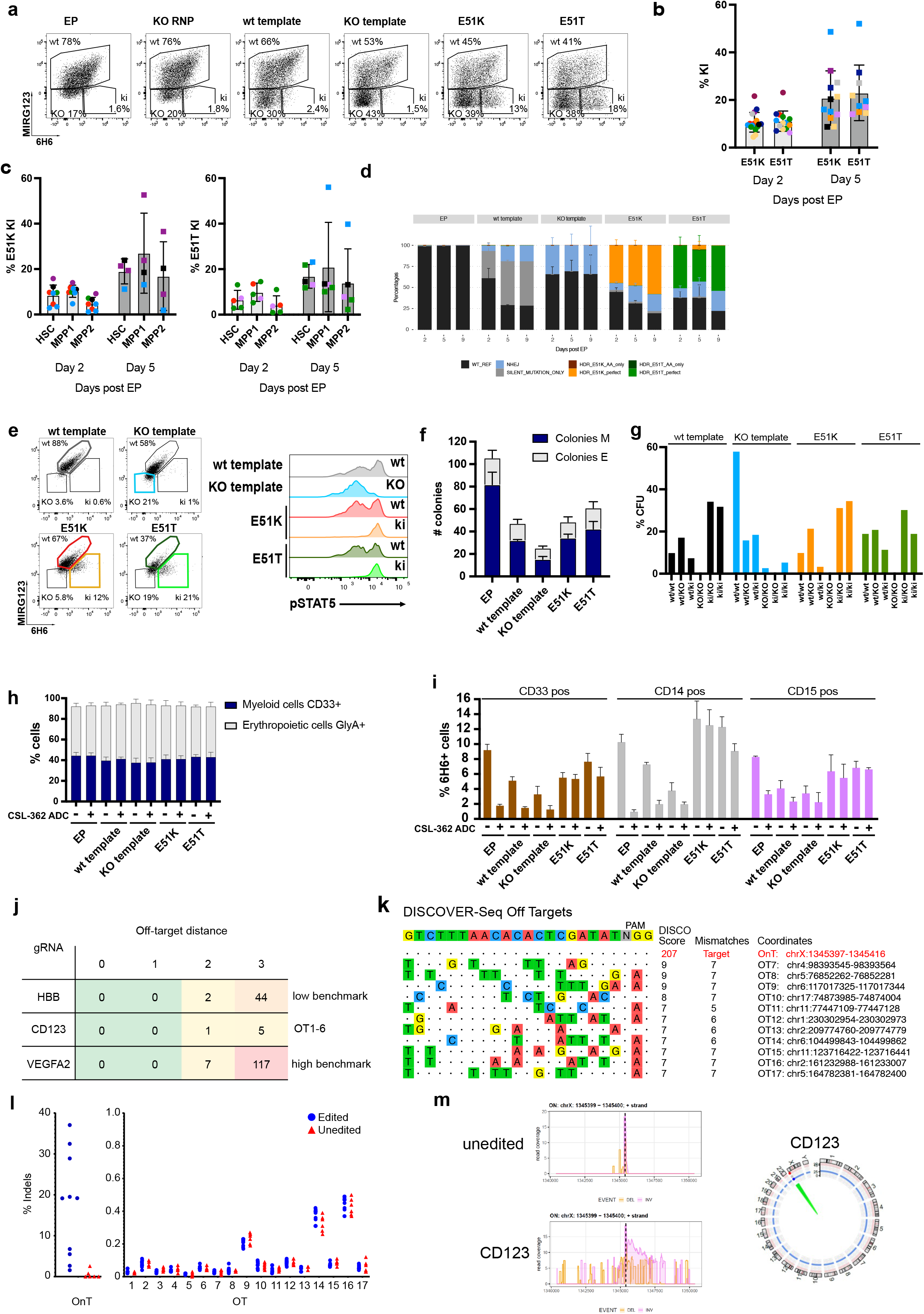
HSPCs expressing CD123 variants E51K and E51T are functional, differentiate normally in vitro and display a good safety profile. **a-m,** Characterization of non-virally CRISPR/Cas9-edited human CD34^+^ HSPCs. **a,** Representative flow cytometry plots showing binding (%) of the anti-human CD123 antibody clones 6H6 and MIRG123 to edited CD34^+^ HSPCs five days after electroporation. EP (cells electroporated with Cas9 protein only), KO RNP (electroporation with RNP only), wt template, KO template, and CD123 E51K or E51T variants (electroporated with respective HDRT). **b,** Frequency of KI cells (MIRG123^-^6H6^+^) two and five days after electroporation. Data from 8 individual donors (each a color) performed in 6 independent experiments with 2 to 4 technical replicates. **c,** Quantification of the KI population in long-term repopulating HSCs (LT-HSC; CD34^+^CD38^-^CD90^+^ CD45RA^-^), multipotent progenitor 1 (MPP1; CD34^+^CD38^-^CD90^-^CD45RA^-^) and MPP2 (CD34^+^CD38^-^CD90^-^ CD45RA^+^). **d,** Representative Amplicon-NGS sequencing of the targeted CD123 locus at 2, 5 and 9 days post-editing of control (EP), wt template, KO template, E51K and E51T conditions. Data from 4 different experiments performed with different donors were pooled. **e,** Representative FACS plots of CD123 stained with 6H6 and MIRG123 (left) and histograms of phosphorylated STAT5 (right) upon exposure to IL-3 in non-virally edited HSPCs. Color-coding in FACS plots and histogram is identical. Data represents 4 independent experiments. **f,** In vitro differentiation of CD123-engineered HSPCs assessed by number of Colony Forming Units (erythroid: E, myeloid: M). CFU were scored using STEMVision based on morphological characteristics. **g,** Allele frequency of the CD123-engineered HSPCs in minimum of 38 colonies. **h,** Frequency of GlycophorinA (GlyA)^+^ and CD33^+^ non-virally edited HSPCs cultured in high cytokine medium with or without CSL-362-ADC for fourteen days. Myeloid lineage: CD33^+^ Erythroid lineage: GlyA^+^. Data from three experiments performed in triplicates. **i,** Frequency of 6H6^+^ CD123-positive cells in the CD33^+^, CD14^+^ or CD15^+^ subsets. **j,** Computational off-target prediction. HBB and VEGFA2 as benchmarking gRNAs. On-target (OnT), Off-target (OT1-6). **k,** DISCOVER-Seq in KO RNP edited HSPCs. On-target (OnT), Off-target (OT7-17). **l,** rhAMPSeq validation of computational prediction and DISCOVER-Seq analysis. Shown are the editing rates as percentage of indels detected at the on-target (OnT) site and at the 17 off-target (OT) sites. Blue circles: Edited category comprises samples treated with gRNA for CD123 (KO RNP, KO template, E151K and E15T). Red triangles: Unedited category comprises samples not treated with gRNA for CD123 (HSC, EP). Samples were obtained from 5 different experiments and donors. **m,** CAST-Seq in unedited and KO RNP edited HSPCs. Coverage plots of the on-target site in *CD123* indicate large inversions (pink line) and deletions (orange line), respectively. Circos plot is used to illustrate chromosomal translocations.

As an orthogonal engineering approach, we used AAV6-mediated HDR. Consistent with the data from the non-virally engineered HSPCs, IL-3 induced a strong pSTAT5 signal in both control HSPCs (non-edited and CCR5 edited) as well as E51K and E51T KI HSPCs (**Extended Fig. 3f, related to Fig. 5e**). In contrast, CD123 KO cells displayed strongly reduced pSTAT5 signalling. As with the non-virally engineered HSPCs, the relative distribution of colonies representing different lineages (BFU-E & CFU-E, CFU-G/GM and CFU-GEMM) was comparable among all genotypes independent of IL-3 (**Extended Fig. 3g, related to Fig. 5h**. Thus, E51K and E51T variants were efficiently engineered into HSPCs using two orthogonal approaches, resulted in functional CD123 receptor expression with intact IL-3 signalling and normal in vitro differentiation capacity yet were shielded from a CD123-targeted ADC.

To assess the safety of the gRNA used to engineer the E51K and E51T variants we used computational prediction, Discover-Seq^41^ and CAST-Seq^42,43^. We used Cas-OFFinder to identify potential off-target sites linked to NGG PAM sequences^44^. Considering sites with a maximum of three mismatches or two mismatches with one DNA and/or RNA bulge identified six candidate off-target sites (OT1-6) (**Fig. 5j and Extended data table 1**). Using these parameters, the gRNA targeting VEGFA resulted in a high number of predicted off-targets as previously reported (high benchmark)^45^. In contrast and in line with previous reports, the gRNA targeting the HBB locus had a low number of predicted off-targets (low benchmark)^41,46^. Compared to these benchmarking gRNAs, the gRNA targeting CD123 had very low numbers of predicted off-targets. Next, we performed Discover-Seq and obtained a typical on-target profile at a good sequencing depth resulting in a Disco score of 207. In contrast, all nominated off-target hits had a very low Disco score (≤9) and at least 5 mismatches. Since Disco scores highly correlate with eventual indel frequencies^41^, these results suggested that the nominated off-target hits were likely false positives (**Fig. 5k and Extended data table 2)**. We selected the top 11 nominations for validation by rhAmpSeq and classified all other hits as false positives. Multiplexed amplicon-NGS on the selected 18 loci (1 on target (OnT), 6 computational, 11 Discover-Seq) performed on unedited HSPC and EP (“unedited”) and KO template, KO RNP, E51K and E51T samples (“edited”) from 4 different donors did not show any evidence of indels (**Fig. 5l**). Finally, as an additional orthogonal assay we performed CAST-Seq analysis, which identifies chromosomal translocations and on-target site aberrations, and nominates off-target sites. An aliquot of the same samples used for Discover-Seq was kept in culture for 5 days post electroporation. In samples edited with RNP complexes, CAST-Seq revealed chromosomal aberrations at the on-target site, such as large deletions and inversions, whereas unedited samples only showcased background activity (**Fig. 5m, left panel**). At the same time, off-target mediated translocations could not be detected (**Fig. 5m, right panel**), confirming the high specificity of the used CRISPR-Cas nucleases. Thus, the selected gRNA did not result in any detectable off-target activity in all the assays performed, demonstrating safe engineering of HSPCs.

Next, we tested in vivo engraftment and differentiation potential. We injected the non-virally engineered HSPCs (EP, wt template, KO template, E51K and E51T) into immunodeficient NBSGW mice. After 16 weeks, all mice of the different groups showed comparable engraftment of hCD45^+^ cells in bone marrow (**Fig. 6a)** and peripheral blood **(Extended data Fig. 4a**). This was also reflected by a comparable frequency and number of human HSCs in the bone marrow (**Fig. 6b**). Furthermore, all groups displayed multi-lineage differentiation in spleen (**Fig. 6c**). Next, we analysed the presence of KI cells among haematopoietic cells in bone marrow. hCD45^+^ cells expressing wt CD123 cells were found with comparable frequencies in all groups. In contrast, E51K and E51T were only found in mice that received KI HSPCs (**Fig. 6d**). We then used the ability to discriminate KI cells from unedited cells to analyse cells with strong CD123 expression. Therefore, we analysed CD123 on pDCs in spleen (gating strategy **Extended data Fig. 4b**). Although pDC numbers were low, the typical MIRG123^-^6H6^+^ KI cells were only present in the mice injected with E51K or E51T HSPCs (**Fig. 6e, f**). Thus, engineered HSPCs harbouring single amino acid substitutions engrafted in vivo and differentiated into multiple cell types including pDCs.

**Fig. 6.**
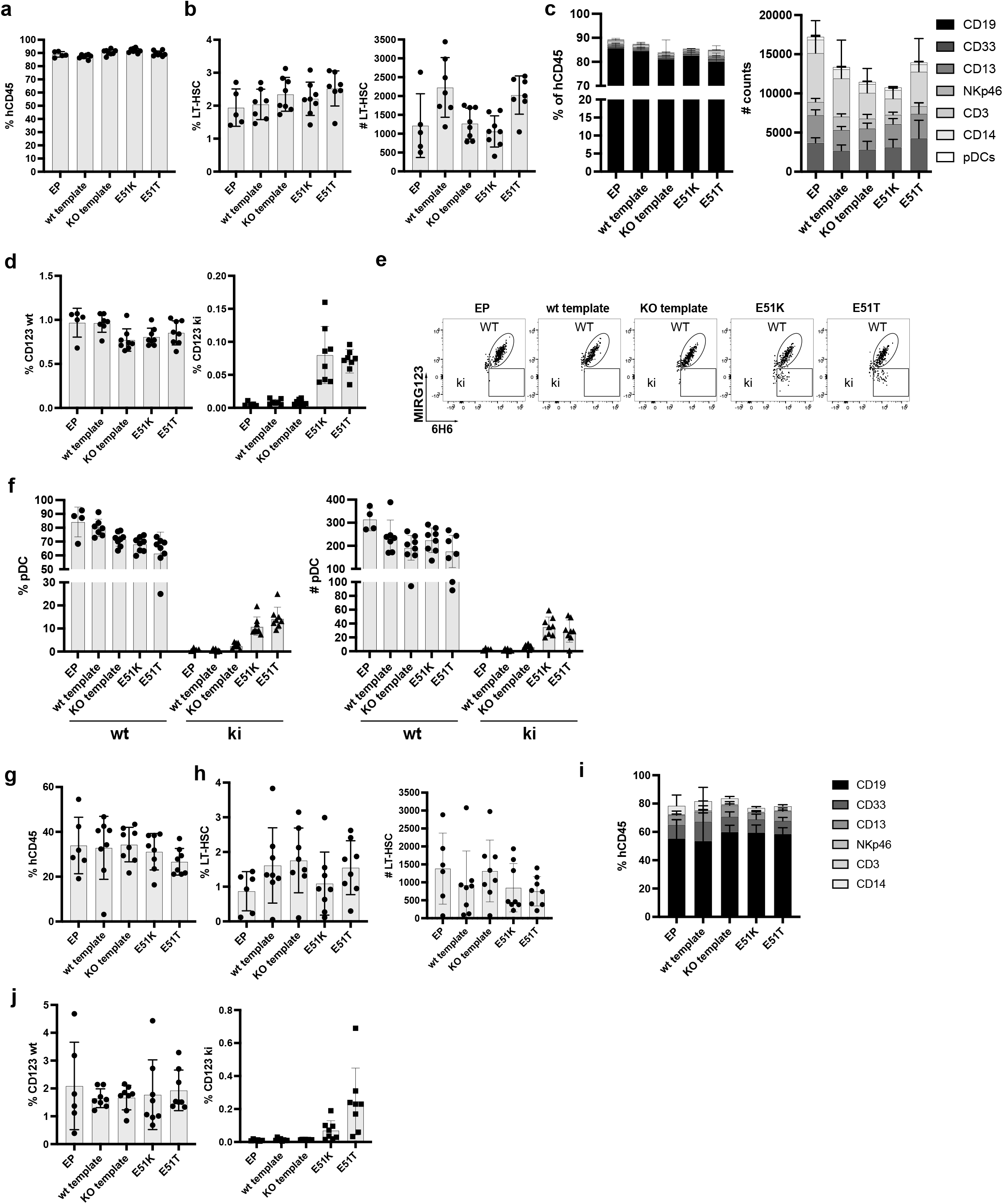
CD123 epitope engineered HSPCs engraft, differentiate normally and possess longterm reconstitution potential in vivo. **a-f,** In vivo engraftment and differentiation potential of non-virally engineered HSPCs expressing E51K and E51T variants measured 16 weeks after injection in NBSGW mice. **a,** Human chimerism (% hCD45^+^) in bone marrow. **b,** Proportion and absolute number of CD34^+^CD38^-^CD90^+^CD45RA^-^ HSCs in the bone marrow. **c**, Multi-lineage differentiation in spleen. Proportions and absolute numbers are shown. **d,** Proportion of wt CD123 (MIRG123+ and 6H6+) **(left)** or CD123 ki (MIRG123- and 6H6+) **(right)** gated on human CD45. **e,** Representative dot plot of pDCs stained with MIRG123 and 6H6 in spleen. Gating strategy to identify pDCs depicted in Extended data Fig.4. **f,** Proportion and absolute number of pDCs. **g-j**, Secondary transplant into NSG-SGM3 mice. **g,** Human chimerism (% hCD45^+^) in bone marrow. **h,** Proportion and absolute number of CD34^+^CD38^-^CD90^+^CD45RA^-^ HSCs in the bone marrow. **i**, Multi-lineage differentiation in spleen. **l,** Proportion of wt CD123 (MIRG123+ and 6H6+) **(left)** or CD123 ki (MIRG123- and 6H6+) **(right)** gated on human CD45. **a-j,** Error bars: mean ± SD.

To investigate whether LT-HSCs were correctly edited and functional we performed secondary transplants. Secondary host mice (NSG-SGM3) transplanted with BM cells from any of the 5 groups demonstrated comparable engraftment (**Fig. 6g**) and frequency and numbers of LT-HSCs in bone marrow (**Fig. 6h**). Furthermore, E51K and E51T engineered HSPCs gave rise to multilineage differentiation comparable to the wt template control mice in spleen (**Fig. 6i**). Importantly, MIRG123^-^6H6^+^ KI cells were detectable in mice reconstituted with E51K and E51T HSPCs, respectively but not from any of the control HSPCs (**Fig. 6j**). Thus, these data demonstrate that E51K and E51T were engineered into LT-HSCs that retained their engraftment potential and KI cells persisted longterm.

After establishing efficient engineering, a favourable safety profile and demonstration of preserved function and longterm engraftment potential of E51K and E51T HSPCs, we sought to investigate tumour selective CD123-targeted immunotherapy and resistance of engineered HSPCs. We co-cultured engineered HSPCs with control T cells not expressing a CAR (CD3^+^ CAR neg) or 123CAR (GFP^+^) together with the AML cell line MOLM-14 (mCherry^+^). Tumor cells were cleared in all samples with 123CAR but not CAR negative control T cells (**Fig. 7a**). Quantification of HSPCs based on absolute cell counts revealed that wt template CD34^+^ HSPCs were depleted in comparison to KO HSPCs and E51K and E51T HSPCs (**Fig. 7b**). Since HSPC KO/KI engineering usually remained < 40% (**Fig. 5a-d**), a 2-fold reduction of CD34^+^ HSPCs would be expected due to killing of unedited, CD123 expressing HSPCs. To verify which cells were specifically eliminated we compared the staining profile of CD123 on the remaining CD34^+^ HSPCs after co-culture. We stained CD123 using only mAb clone 6H6 to avoid possible epitope masking. HSPCs of all 4 groups incubated with CAR negative control T cells expressed CD123 (**Fig. 7c**). In contrast, 123CAR eliminated CD123 expressing cells in wt template HSPCs and KO template HSPCs but not E51K or E51T HSPCs. Thus, 123CAR preferentially depleted highly CD123 expressing HSPCs. However, quantification showed that wt template HSPCs were also depleted among the 6H6^-^ cells since very few cells remained (**Fig. 7c, d**). In contrast, E51K and E51T HSPCs were resistant and continued to express CD123. In a similar experiment we incubated MOLM-14 cells (CTV) with non-engineered HSPCs (EP) and 123CAR cells in which case we observed near complete elimination of all EP control HSPCs while E51K and E51T HSPCs were resistant (**Extended data Fig. 5a, b**). Together, these results suggest 123CAR-mediated bystander killing of HSPCs that express low or no CD123. 123CARs also efficiently depleted other AML cell lines (OCI-AML2, OCI-AML3) as well as patient derived AML cells (PDX) (**Extended data Fig. 5c**). Since PDX cells were efficiently killed by 123CAR, we next mixed PDX (CTV^+^) with edited HSPCs and 123CAR. As observed for MOLM-14, 123CARs indiscriminately killed PDX as well as wt template HSPCs (**Fig. 7e, f**). In contrast, 123CAR displayed selective cytotoxicity towards PDX cells but preserved E51K and E51T HSPCs (**Fig. 7e-h**).

**Figure 7.**
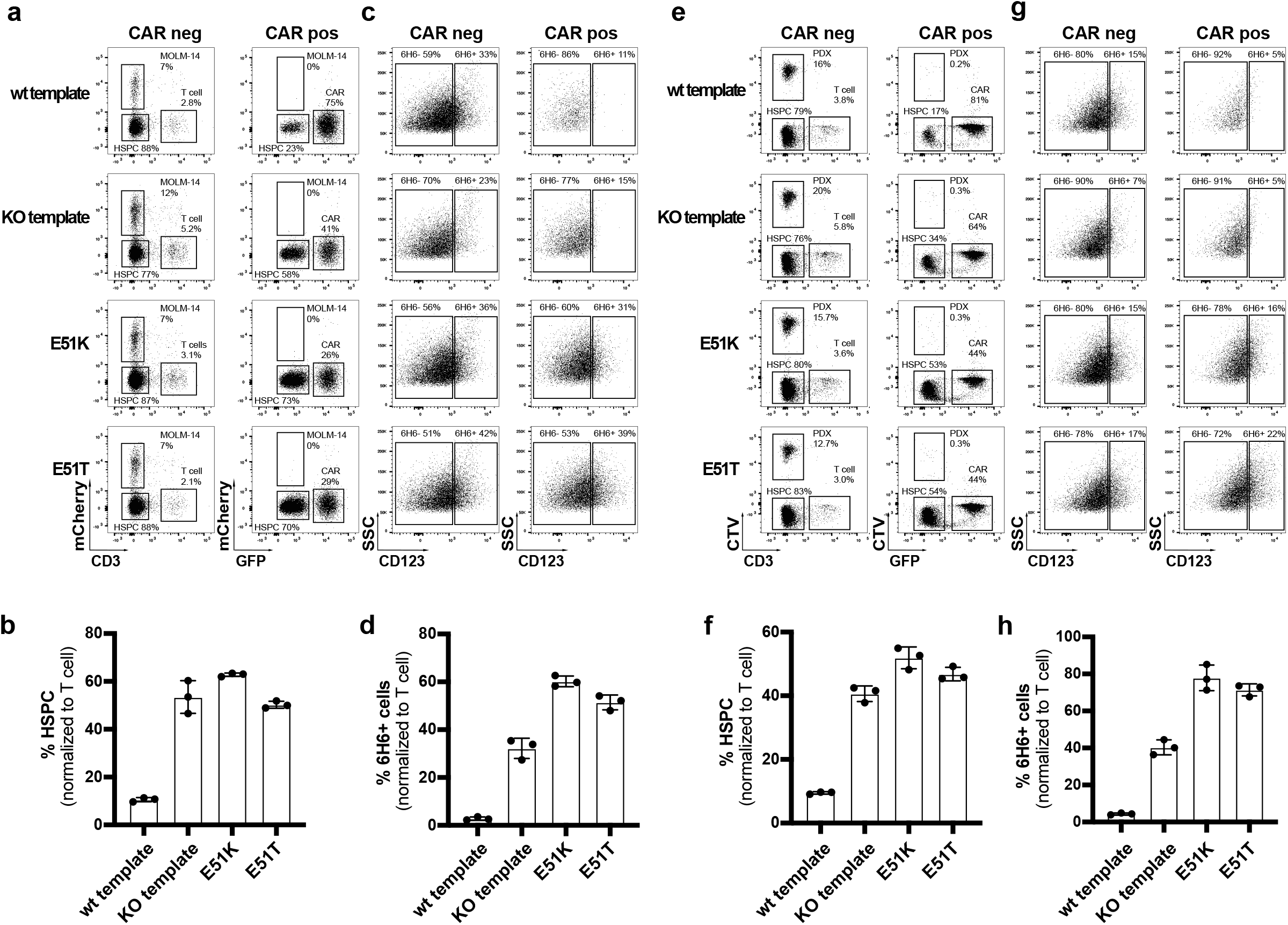
Engineered HSPCs enable tumor-selective CD123 immunotherapy. **a-d,** Non-virally edited HSPCs co-cultured with MOLM-14-mCherry (AML cells) and control T cells or 123CAR for 2 days. **a,** Representative dot plots indicating proportion (%) of CD3^+^T cells, MOLM-14 cells and non-virally edited CD34^+^ HSPCs on day 2 of co-culture. **b,** Quantification of **(a)** based on absolute counts. **c**, FACS plots illustrating proportion (%) of wt or ki HSPCs at the end of the co-culture based on the binding characteristics to the mAb 6H6. Only clone 6H6 was used to avoid epitope masking by the 123CAR. **d,** Quantification of **(c)** based on absolute counts. **e-h,** Non-virally edited HSPCs co-cultured with PDX (CTV labeled) and control T cells or 123CAR for 2 days. **e,** Representative dot plot indicating proportion (%) of CD3^+^T cells, PDX cells and non-virally edited CD34^+^ HSPCs on day 2 of co-culture. **f,** Quantification of **(e)** based on absolute counts. **g**, FACS plots illustrating proportion (%) of wt or ki HSPCs at the end of the co-culture based on the binding characteristics to the mAb 6H6. **h,** Quantification of **(g)** based on absolute counts. Data normalized to co-culture with T cells. **a-h,** Error bars: mean ± SD.

Next, we investigated selective cancer immunotherapy by the TCE and ADC. The CSL362/OKT3-TCE was less potent than the 123CAR since some MOLM-14 cells remained (**Extended data Fig. 5d**). Nevertheless, we also observed an overall depletion of EP HSPCs suggesting some bystander killing of HSPCs expressing low or no CD123 (**Extended data Fig. 5d, e**). To exclude possible artifacts due to epitope masking by remaining CSL362/OKT3-TCE that could block MIRG123 binding, we performed an experiment without MOLM-14 cells and FACS-sorted HSPCs based on 6H6 staining on the day of analysis. Sanger sequencing confirmed that EP HSPCs (6H6^-^ and 6H6^+^) and 6H6^-^ cells sorted from E51K and E51T HSPCs were primarily WT (**Extended data Fig. 5f**). Thus, these are HSPCs that have a CD123 WT genotype but do not express CD123. In contrast, sorted 6H6^+^ KI HSPCs were almost purely E51K (AAG) or E51T (ACC), respectively. Thus, E51K and E51T expressing HSPCs withstood CD123-targeted TCE-mediated cytotoxicity leading to an enrichment of edited KI HSPCs. To test ADC as a depletion modality we tested selective tumour killing with MOLM-14 and engineered HSPCs. The CD123-ADC depleted CD123 expressing 6H6^+^ HSPCs but not E51K or E51T HSPCs (**Extended data Fig. 5h**). Thus, engineered KI HSPCs were also shielded from an ADC. However, MOLM-14 cells were not depleted and HSPC numbers for EP, E51K and E51T remained the same (**Extended data Fig. 5g**). Thus, we did not observe any bystander killing by this CD123-targeted ADC but overall efficacy was limited. In summary, epitope engineered, shielded HSPCs can safely be generated, are functional but resistant to CD123-targeted immunotherapy and thereby enable tumour-selective targeting.

## Discussion

HSCT is a potentially curative approach for haematologic diseases. A conditioning phase prepares the transplantation of autologous or allogeneic HSCs which, after engraftment, will rebuild a new haematopoietic and immune system. The conditioning serves to remove host HSCs and can kill tumour cells when HSCT is applied for malignant diseases. However, current untargeted cytotoxic conditioning regimens have been directly or indirectly associated with transplant related morbidity and mortality. With the advent of highly effective targeted depleting agents such as mAbs, ADCs, TCEs and CAR T cells it may become possible to replace untargeted conditioning and tumour control by antigen-specific immunotherapy^25,47–50^. However, the absence of suitable antigens constitutes a major impediment to progress in this field^11,12,22^. In particular, targeting LSCs would be highly desirable to treat AML but the risk for myelosuppression arising from shared antigen expression of LSCs and healthy HSCs precludes a continuous posttransplant therapy^25^. As a compromise, it was proposed to use CAR T cells directed against CD123 or CD117 to purge tumour cells and HSCs alike, followed by myeloablative conditioning to remove the CAR T cells before HSCT^21,24,25^. Recently, CAR T cells targeting CD7 were used clinically to deeply purge relapsed T cell acute lymphoblastic leukaemia^51^. However, the CAR7 were only used for a limited time before HSCT and they led to multilineage cytopenia. Thus, the targeted immunotherapy was limited to the pretransplant period and only served as a bridge-to-transplant. However, the ability to continue immunotherapy posttransplant would be highly desirable to eliminate MRD and prevent relapse longterm.

We previously used genome engineering to substitute single amino acids in surface proteins of murine T cells which completely abolished binding of specific mAbs^52^. Since we converted protein variants known to have exchangeable function (aka congenic markers) the results suggested that engineering a minimal alteration in surface proteins could enable safe cell shielding for therapeutic purposes. To enable HSCT with continuous antigen-specific immunotherapy, we engineered human HSPCs to express epitope engineered CD123 and demonstrate that these cells are both shielded and functional, respectively. We used structural information to identify at least one aa substitution that completely abrogated mAb binding for 3 separate residues. This is important since it increases the chances that an appropriate genome engineering strategy can be found to safely and efficiently insert a variant into primary therapeutic cells. The choice of the most suitable genome engineering approach will depend on the desired aa substitution and cell type. To establish feasibility, we employed CRISPR/Cas9-mediated HDR which provides flexibility and precise programmability through a DNA template encoding the desired amino acid exchange. Despite some shortcomings such as cellular toxicity, HDR-mediated HSPC editing is safe and clinically relevant^53,54^. In fact, we did not find any detectable off-target activity using multiple complementary assays, demonstrating safe engineering of HSPCs. However, for clinical translation additional studies will be needed, e.g. tumorigenicity studies. In addition, more recently developed genome engineering approaches provide additional options. Specifically, base editing is well tolerated, can achieve high editing efficiencies in HSPCs and is suitable for multiplexed editing^55^. However, base editors have different constraints than HDR since they typically only convert A→;G or C→;T, also have PAM restrictions and the editing window - despite being limited - can result in unwanted bystander editing. As a consequence, base editing can only install a fraction of desired codon changes for a particular aa change. Indeed, we found that these inherent limitations are relevant when using base editors to install cell shielding variants because the most suitable amino acid substitutions may not be accessible to base editing or bystander editing may affect the function of the engineered protein (Marone & Jeker, unpublished observations). Therefore, careful characterization of the editing outcome and the resulting protein variants is required (Garaudé & Jeker, unpublished observations). Prime editing employs a templated approach that also avoids dsDNA breaks and provides the greatest flexibility to install any desired codon change and could therefore be suitable for epitope engineering^55^. However, prime editing was only recently shown to work in HPSCs and further investigation is needed^56^.

Information about naturally occurring polymorphisms without known disease association (e.g. CD123 E51K) could inform variant choice from a safety perspective. For instance, although IL-3 binding to CD123 E51K may be slightly reduced compared to WT CD123 (**Fig. 4d**), it appears that different binding strengths to IL-3 may be biologically tolerated. Theoretically, allogeneic HSPC donors could be pre-screened for such variants which would provide shielding without the need to engineer the cells. However, practically, most polymorphisms are likely too rare since HSPCs also have to be matched for HLA. Conversely, this indicates that some patients receiving therapeutic cell depleting agents may be non-responders due to genetic variation. Remarkably, our results show that a single aa substitution is sufficient to protect from mAb (ADCC), ADC, TCE and even CAR T cells. Thus, shielded HSPCs or other therapeutic cells could be combined with a broad range of available cell depleting agents. However, our data shows that careful characterization of the variants for potential clinical translation is important. For instance, although R84E shielded from all tested depletion modalities, it showed signs of reduced stability. Furthermore, S59P and R84E both showed reduced IL-3 binding and therefore were not further pursued. In addition, the degree of binding reduction will be relevant for the choice of the MoA of the depleter. For instance, S59Y, R84T and R84Q protected from ADCC and TCE but led to substantial killing by the CAR T cells. It is noteworthy that the CAR T response was rather digital with non-binders being protected whereas weak binders resulted in strong CAR T activation and cytotoxicity. In contrast, responses to weak binders were more analogue for ADCC and TCE resulting in much more limited killing. The sensitivity of CAR T responses even to weak binding will also be important when considering targets with expression by non-hematopoietic cells. For instance, although Flt3 could be an attractive target to treat AML, Flt3 inhibitors may lead to acute cardiotoxicity^57^. Flt3 expression was found in murine cardiomyocytes and expression in human iPS-derived cardiomyocytes was sufficient to result in mild but detectable activation of Flt3-targeted CAR T cells^58,59^. Thus, the therapeutic window created by a shielding variant will be a function of the binding reduction between the WT and the engineered variant, the expression on the target cells (which may be dynamic), expression on non-hematopoietic cells as well as the efficacy of the depleting agent. In some cases, e.g. when a blocking Ab is used, residual binding of the shielding variant may be acceptable. In others, when the variant is to be paired with a highly effective depleter (e.g. ADC or CAR T), the non-binding must be more stringent.

In order to fully exploit the advantage of combining shielded HSPCs with targeted immunotherapy, the function of the antigen should be preserved^11^. Our results show that CD123-deficient HSPCs had a competitive disadvantage in vitro and CD123 KO HSPCs were strongly depleted. In contrast, selected shielded CD123 single amino acid substitution variants preserved IL-3 binding, signalling (p-STAT5) and IL-3 dose-dependent growth. Furthermore, engineered KI HSPCs differentiated normally in vitro, engrafted and displayed multi-lineage differentiation potential in vivo. Secondary transplantation experiments confirmed successful engineering of longterm reconstituting LT-HSCs. Thus, molecularly shielded HSPCs could allow tumour-selective CD123 targeted immunotherapy and in parallel enable rebuilding a CD123 variant-expressing haematopoietic system. For instance, CD123 targeting therapies are clinically successful for the treatment of BPDCN, a pDC-derived neoplasm. However, also healthy pDCs strongly express CD123 and therefore will be depleted by any effective CD123 immunotherapy. Yet, pDCs are primary producers of type I interferon and hence anti-viral immunity. Their importance was recently illustrated for the protective immunity against SARS-CoV-2 ^60^. Therefore, transplanting CD123-shielded HSPCs may in the future potentially allow to fully restore immunity with a completely functional haematopoietic system while simultaneously allowing efficient tumour immunotherapy. More broadly, we envision that function-preserving but shielding variants could be identified for other proteins that currently cannot safely be targeted. Applications could include T cell malignancies since T cell deficiency results in unacceptably severe immunosuppression and T cell function cannot readily be replaced. CD7 is of interest to treat T cell malignancies and AML but CAR7 are currently only used as a bridge-to-transplant^51^. Importantly, preserving the function of the engineered proteins could enable multiplex shielded HSPCs that are resistant to combination immunotherapies to address disease heterogeneity. However, safety of multiplex edited HSPCs would need to be assessed carefully. In addition, directed ablation of HSPCs or more generally haematopoiesis has been a long-standing goal to improve HSCT^11^ and could be applied beyond malignancies. CD117 and CD45 are promising targets but current approaches are limited to short-term depletion^24,25,47–50^. Targeting CD45 could be applied to systemic autoimmune diseases but it is currently unknown whether short-term depletion will be sufficient and clinically tolerated^48,61,62^. Shielded CD45 variants could enable a gradual replacement of the haematopoietic system including autoreactive lymphocytes. Finally, epitope shielding could open up development of new immunotherapies against as yet unidentified targets and may be applicable for replacement of other cell types e.g. T, B and the many other immune cells which are currently being considered for engineered cellular therapies^11^.

## Acknowledgements

We acknowledge the contributions of the following research core facilities at the University of Basel and the Department of Biomedicine: the teams of the animal facility for expert animal husbandry and the flow cytometry core facility for excellent support. We thank Marko Hasiuk for gRNA design; Jonathan Knowles for critical reading of the manuscript; Dario Neri and Mihaela Zavolan for sharing reagents; Marko Hasiuk, Simon Garaudé and Hanna Studer for experimental support; Andreas Zingg for advice for CAR T experiments. Some figures created with BioRender.com. This project has received funding from the European Research Council (ERC) under the European Union’s Horizon 2020 research and innovation programme (grant agreement No. 818806 to L.T.J.). J.E.C. and D.G. are supported by the NOMIS Foundation, the Lotte and Adolf Hotz-Sprenger Stiftung, SNSF grant 310030_188858, and the Helen Hay Whitney Foundation. T.Ca. and T.I.C. acknowledge funding from the German Federal Ministry of Education and Research (BMBF) within the Medical Informatics Funding Scheme - EkoEstMed–FKZ 01ZZ2015 (G.A.).

## Author contributions

These authors contributed equally: R.M., E.L., A.D., R.L.; E.L., A.D., R.M. designed and performed most experiments, analysed data and wrote the manuscript; R.L. designed and performed computational analysis and wrote the manuscript; C.E., A.W., L.G.P., L.X. performed experiments and analysed data; G.C. performed experiments and produced the TCE; F.S. generated and provided MOLM-14-mCherry-luciferase cells; C.L. and J.T.B. generated and provided AML PDX samples; A.DA. performed in vivo experiments supervised by R.M.; J.Z. and T.B. performed and analysed rhAmpSeq; D.S. analysed NGS data; A.S., A.H., A.C. performed protein stability experiments; V.DS., C.D. performed HSPC isolation; D.G., M.S. performed Discover-Seq and bioinformatics analysis under supervision of J.E.C; J.R., M.R., G.A. performed CAST-Seq and bioinformatics analysis under supervision of T.I.C. and T.CA.; L.B. performed initial computational analysis; T.S. supervised computational analysis; M.P. supervised AAV experiments; S.U. designed and supervised experiments; L.T.J. conceived the study, supervised experiments, acquired funding and wrote the manuscript. All authors read and approved the manuscript.

## Data availability Statement

Datasets are available from the corresponding author on reasonable request.

## Materials and Methods

All antibody sequences, gRNAs, oligos and flow cytometry antibodies used in the study are listed in Extended data table 4.

### Structural dataset and computational design of CD123 protein variants

Experimentally determined three-dimensional structures of the CD123 protein were retrieved from the PDB and include (i) the CD123-CSL362 complex in open and closed conformation (PDB ID: 4JZJ^1^), (ii) the CD123-IL-3 binary complex (PDB ID: 5UV8^2^), (iii) the CD123 unbound structure extracted from the complex in open and closed conformation. Given the structure of the complexes, the CSL362 and IL-3 epitopes were defined as the sets of CD123 residues having at least one atom within 4Å to any CSL362 and IL-3 atoms, respectively. Per-residue relative solvent accessibility area was computed using the Lee & Richards algorithm^3^ implemented in FreeSASA^4^, using default parameters and upon removing crystallographic waters, sugars and ions. Comprehensive mutagenesis was performed *in silico* using the EVmutation sequence-specific probabilistic model^5^ and the mutational effect, defined as statistical energy difference ΔE, estimated using the epistatic coevolutionary analysis framework^6s8^. For a given mutant, the ΔE is computed as the sum of differences of the constraints on individual amino acid sites plus the sum of differences of the coupling parameters computed for all pairs of sites involving the mutated site. In order to quantify the total epistatic constraint acting on a given amino acid site of interest, evolutionary coupling analysis was run on a multiple sequence alignment (MSA) spanning the entire CD123 sequence. The MSA was built using five iterations of the jackhammer HMM search algorithm against the non-redundant UniProtKB database^9^ and default significance score for inclusion of homologous sequences. As values of ΔE below, equal and above 0 correspond to putatively beneficial, neutral, and deleterious effects, respectively, variants at a given site were selected upon ranking for decreasing ΔE.

### Eukaryotic cell lines

Freestyle CHO-S cells were purchased from Thermo Scientific (Cat#R80007) and were expanded in PowerCHO 2 Serum-free Medium (Lonza BELN12-771Q) supplemented with GlutaMAX (Gibco), HT supplement (Cat# 41065012) and antibiotic-antimycotic (Cat#15240062), to a maximum density of 20×10^6^ cells/ml. TF-1 were purchased from DSMZ (Cat#ACC334) and maintained in RPMI-1640 media supplemented with 10% heat-inactivated FCS (Gibco Life Technologies), 2mM GlutaMAX and 2ng/ml hGM-CSF (215-GM, Bio-Techne). HEK-293 cells were a kind gift of M. Zavolan (Biozentrum Basel) and cultured in Dulbecco’s Modified Eagle’s Medium - high glucose (Sigma-Aldrich) supplemented with 10% heat-inactivated FCS and 2mM GlutaMAX. MOLM-14 were purchased from DSMZ (Cat#ACC777) and K562 were purchased from ATCC (Cat#CCL-243). Both cell lines were maintained in RPMI-1640 media supplemented with 10% heat-inactivated FCS (Gibco Life Technologies) and 2mM GlutaMAX. All cell lines were freshly thawed and passaged 3-6 times prior to use. AML cell lines (MOLM-14, OCI-AML-2 and OCI-AML-3) were retrovirally transduced with MI-Luciferase-IRES-mCherry (gift from Xiaoping Sun, Addgene plasmid #75020; http://n2t.net/addgene:75020; RRID:Addgene_75020). Cells were then FACS-sorted based on mCherry expression. After expansion, cells were STR profiled and tested mycoplasma negative before being frozen until further use.

### Hematopoietic cells from human subjects (PDX)

De-identified patient-derived AML samples were obtained from PDX repository (Cancer Research Center of Toulouse, France)^63,64^. Cells were propagated in NSG mice for 2 generations. A signed written informed consent for research use in accordance with the Declaration of Helsinki was obtained from patients.

### Cloning and expression of recombinant wildtype human CD123 and its variants

Full length cDNA of human CD123 (NM_002183.2) was obtained from a pCMV3 vector (Cat#HG10518-M, Sino Biological). The Hygromycin sequence was replaced by a Neomycin resistance cassette by Gibson Assembly. The point mutations of the human CD123 variants were introduced into the vector using PCR (see Extended data table). 2×10^6^ HEK-293 cells were electroporated with the pCMV3 vector encoding CD123 wildtype or its variants using the Neon Transfection System (ThermoFisher; 1100V, 20ms, 2pulses). To generate stable cell lines Geneticin G418 (50mg/ml; BioConcept) was added to the cell culture medium at a concentration of 350µg/ml.

### Cloning and expression of the monoclonal antibody CSL362 biosimilar (MIRG123)

The heavy and kappa light chain variable regions (VH and VKL) of CSL362 were derived from the CSL362/OKT3-TCE^10^. To generate the monoclonal IgG1 antibody CSL362 biosimilar MIRG123 the VH and VKL sequences were cloned into AbVec2.0-IGHG1 (Addgene plasmid # 80795) and AbVec1.1-IGKC (Addgene plasmid # 80796), respectively, kindly provided by Hedda Wardemann^11^. 2×10^9^ Freestyle CHO-S cells were resuspended in 500ml ProCHO 4 Protein-free Medium (Lonza, Cat#BEBP12-029; supplemented with 1XHT, GlutaMAX, antibiotic-antimycotic) and co-transfected with both plasmids (0.6mg each) using 5mg polyethyleneimine (PEI, Polysciences, Cat#23966). The cells were expanded under constant rolling (140rpm) at 31°C, 5% CO2 for 6 days. The Freestyle CHO-S cells were pelleted and the filtered supernatant (0.22μm filter) was applied on a Protein A column for purification. CSL362 biosimilar was eluted with 0.1M glycine (pH=2.2), 0.5ml fractions were collected and OD280 was measured using a Nanodrop spectrophotometer. The high-concentration fractions were pooled and dialyzed twice overnight in PBS.

### Expression and purification of CSL362/OKT3-TCE

The design of the CSL362/OKT3-TCE was recently described^10^ and the plasmid was a kind gift from D. Neri (ETH Zurich). Freestyle CHO-S producer cells were transfected using PEI (as described above) with 1.7mg TCE DNA. Following 6 days of expansion the filtered cell culture supernatant was applied to a 5ml Ni-NTA column (ThermoFisher) prewashed with 100ml washing solution (PBS, 150mM NaCl, 5mM Imidazole, pH7.4) for protein purification. The eluted and dialysed TCE was filtered (0.22μm) and stored in aliquots (1mg/ml) at -80°C.

### Flow cytometry and cell sorting

Flow cytometry was performed on BD LSRFortessa with the BD FACSDiva Software, and the data was analysed with FlowJo Software. Antibodies used for flow cytometry can be found in the Extended data table. For cell sorting the cells were pelleted and resuspended in FACS Buffer (PBS + 2%FCS) supplemented with 1mM EDTA, and sorting was performed either on BD FACSAria or BD FACSMelody Cell Sorter. The control cells were also subjected to the sorting process.

### Primary human T cell isolation and culture

Leucocyte Buffy coats from anonymous healthy human donors were purchased from the blood donation centre Basel (Blutspendezentrum SRK beider Basel, BSZ). Peripheral blood mononuclear cells (PBMCs) were isolated by density centrifugation using SepMate tubes (Stemcell technologies) and the density gradient medium Ficoll-Paque (GE Healthcare) according to the manufacturer’s protocol. Human T cells were purified (> 96% purity) by magnetic negative selection using an EasySep Human T Cell Isolation Kit (Cat#17951, Stemcell Technologies) according to the manufacturer’s instruction. If frozen PBMCs were used, T cells were isolated after thawing and cultured in supplemented media without stimulation overnight. T cells were cultured in RPMI-1640 Medium (Sigma-Aldrich) supplemented with 10% heat-inactivated human serum (AB^+^, male; purchased from BSZ Basel), 10mM HEPES (Sigma-Aldrich), 2mM GlutaMAX, 1mM Sodium Pyruvate, 0.05mM 2-Mercaptoethanol, 1% MEM Non-essential amino acids (100x) (all Gibco Life Technologies) and IL-2 150U/ml (Proleukin, University Hospital Basel). The medium and IL-2 was replenished every 2 days, and the cells were kept at a cell density of 1×10^6^ cells/ml.

### ADCC assay (FcγRIIIa activation assay)

ADCC assay was performed using the ADCC Reporter Bioassays, V Variant (Promega, Cat#G7015) according to the manufacturer’s instructions. Target cells HEK-293 expressing the CD123 variants were seeded in white 96 well-plate clear bottom at 4400 cells/100μl culture medium. At day 1, medium was removed and effector cells (Jurkat/FcγRIIIa/NFAT-Luc cells; E:T ratio 12:1) and MIRG123 antibody (final concentration 1μg/ml) were added and incubated for 5h at 37 °C 5% CO2. The emitted luminescence was read 10min after the addition of Bio-Glo Luciferase Assay Reagent (Promega) using the PHERAstart FSX (BMG LABTECH) program Luc-Glo (LUM), GainA=3600, Optic module=LUMplus. Raji cells incubated with 1μg/ml of anti-CD20 antibody rituximab were used as positive control.

### In vitro TCE-mediated killing assay with primary T cells

For the TCE killing assays the HEK-293 target cells were co-cultured together with human effector T cells and the CD3/CSL362-TCE at an effector to target ratio of 10:1 for 72 h. One day prior to co-culture HEK, HEK-CD123 and the HEK expressing the CD123 variants were stained with CellTraceViolet (CTV) according to the manufacturer’s protocol. The following day effector T cells were added with the TCE at a concentration of 300ng/ml and kept for 72h at 37 °C. Cytotoxic activity and activation of T cells were analysed by flow-cytometry. Specific killing was calculated as follows: (1-No. alive target cells with TCE/No. alive target cells without TCE)*100. Cell morphology was assessed with the light microscope Axio Vert.A1 (Zeiss) at 20x magnification.

### Design and production of the CD123CAR HDRT

The CAR T cells were generated by co-electroporation of CRISPR-Cas9 Ribonucleoproteins (RNPs) specific for the *TRAC* locus and a double-stranded DNA HDR template (HDRT). The HDRT encoding a second-generation CD123-specific CAR with the scFv of clone CSL362, the CD8α hinge and transmembrane domain (Gen CD8A ENSG00000153563), the intracellular signalling moieties 4-1BB (Gen TNFRSF9 ENSG00000049249) and CD3ζ (Gen CD247 ENSG00000198821), as well as the fluorescent reporter protein GFP (for full sequence see Extended data table). It is flanked by symmetric arms of homology (300 bp) complementary to the *TRAC* locus Exon 1. The construct was synthesized by GenScript^®^ and the plasmid was used as template for PCR amplification (Kapa Hifi Hotstart Ready Mix, Cat#F-530-L, Roche). The PCR amplicon was purified with NucleoSpin Gel and PCR clean-up kit (Macherey-Nagel) according to the manufacturer’s instruction and the correct size was verified by gel. The HDRT was then condensed to a final concentration of 1µg/µl using vacuum concentration, and stored at -20°C until usage.

### Engineering primary CD123CAR T cells

Protocols for human CRISPR/Cas9-mediated genome engineering are based on *Roth et al.*^12,13^ In short, Cas9 RNPs were freshly generated prior to each electroporation. Thawed crRNA and tracrRNA (purchased from IDT Technologies, at 200µM) were mixed in a 1:1 molar ratio (120pmol each), denatured at 95°C for 5min, and annealed at room temperature (RT) for 10 to 20min to complex a 80µM gRNA solution. Poly-Glutamic Acid (PGA; 15-50 kDa at 100mg/ml; Sigma-Aldrich) was added to the gRNA in a 0.8:1 volume ratio^14^. To complex RNPs, 60pmol recombinant Cas9 (University of California Berkeley at 40µM) was mixed with the gRNA (molar ratio Cas9:gRNA = 1:2) and incubated for 20min at RT in the dark. Prior to electroporation isolated human T cells were activated for 48h with CD3/CD28 Dynabeads (Thermofisher) at a cell to beads ratio 1:1 together with the recombinant human cytokines IL-2 (150U/ml), IL-7 (5ng/ml; R&D Systems) and IL-15 (5ng/ml; R&D systems). Electroporation was performed with the 4D-Nucleofector^™^ system (Lonza) with Program EH-115. Following activation, the T cells were de-beaded using an EasySep^™^ magnet and 1×10^6^ cells were resuspended in 20µl Lonza supplemented P3 electroporation buffer. HDRT (3-4µg) and RNPs (60pmol) were mixed separately and incubated for 5min. The cells were added to the mix and the total volume was transferred to the 16-well Nucleocuvette Strips. Immediately following electroporation 80µl of prewarmed supplemented medium was added to each cuvette and incubated at 37°C. After 20min, the cells were transferred into 48-well culture plates and replenished wit IL-2 500U/ml. Following flow-sorting at day 3 to 5 post-electroporation the cells were expanded for 5 to 6 days until used for the subsequent experiments. Control T cells were electroporated with an incomplete RNP (missing the specific crRNA), otherwise processed as the CAR T cells.

### In vitro human CD123CAR killing assay

The day before the co-culture HEK-293 target cells were stained with CTV and kept in supplemented human medium overnight. The flow-sorted, expanded GFP^+^ CAR T cells and control cells were added to the target cells in an effector to target ratio 10:1 and co-cultured for 24h. Specific killing and T cell activation was measured by flow cytometry. Specific killing was calculated according to the indicated formula: (1-No. alive target cells in co-culture with CAR T cells/No. alive target cells in co-culture with control cells)*100. Using the microscope Axio Vert.A1 (Zeiss) cell morphology was recorded.

### In vitro 123CAR-mediated killing assay of non-virally engineered HSPCs

For the 123CAR killing assays the HSPCs were edited as described above. Two days after electroporation edited HSPCs, MOLM-14-mCherry, OCI-AML2-mCherry, OCI-AML3-mCherry or PDX (labelled with CTV according to the manufacturer’s instruction) were co-cultured with control T cells or 123CAR in a 96 U-bottom plate at an effector to target ratio of 3:1 for 48 h at 37 °C. In a set of experiments MOLM-14-mCherry or PDX (labelled with CTV according to the manufacturer’s instruction) were co-cultured with edited HSPCs and T cells or 123CAR cells at an effector to target to tumour ratio of 3:0.5:0.5 together. Cytotoxic activity (specific killing and elimination of non-edited HSCs) was analysed by flow cytometry.

### Human cytokine measurement (ELISA)

IFNγ was measured from the supernatants of the co-culture experiments (TCE / CAR) using the colorimetric ELISA MAX Standard Set Human IFNγ kit (BioLegend) according to the manufacturer’s instruction. The optical density was read at 450 nm with the microplate reader. A standard curve calculated from standard dilutions was run in duplicates with every experiment.

### Genomic DNA extraction and sequencing from human T cells, eukaryotic cell lines and HSPCs

Genomic DNA was extracted using the QuickExtract^TM^ (Biosearch technologies; QE09050) according to the manufacturer’s instruction. Alternatively, cells were lysed in Tail Lysis Buffer (100 mM Tris [pH 8.5], 5 mM Na-EDTA, 0.2% SDS, 200 mM NaCl) containing Proteinase K (0.1 µg; Sigma-Aldrich) at 56 °C (1000 rpm). The DNA was precipitated with isopropanol (1:1 volume ratio), and washed in 70% ethanol. The genomic DNA concentration was measured with a NanoDrop™ device (Thermo Fisher). PCR was performed using either GoTaq G2 Green Master Mix (Cat#M782B, Promega) or Kapa Hifi Hotstart Ready Mix. Sanger Sequencing was performed at Microsynth AG Switzerland. Sequences were analysed using MegAlign Pro (DNASTAR).

### Next generation sequencing

Genomic DNA was isolated from HSPCs using Quick extraction buffer. Targeted amplicons library was prepared following Illumina’s recommendation using a two-step PCR protocol. Briefly, nested PCRs were performed on each DNA sample using the HiFi KAPA polymerase (Roche). Following Illumina barcoding (Nextera indices, Illumina), PCR samples were pooled, beads purified and quantified using Qubit dsDNA BR (Thermo Fisher). Library was sequenced on an Illumina MiSeq instrument using Illumina MiSeq Reagent Kit v2 Micro (300-cycles) with 50% PhiX spike-in (Illumina). After demultiplexing, each sample was assessed for quality and analysed using CRISPResso v2 ^REF 65^. For each of the samples, we provided the HDR template, the reference sequence, the guide sequence and applied a minimum base quality of Phred 25. We used a custom R script to quantify each allele within a quantification window of 4 nucleotides including one silent mutation followed by the targeted AA.

### Bio-Layer Interferometry (BLI) measurement of binding

All BLI measurements were performed either on an Octet Red96e (ForteBio) or on an Octet R8 (Sartorius). The extracellular domain (ECD) of CD123 WT and variants were produced and purified by Icosagen.

### CSL362 hIgG1 binding to CD123 WT ECD and variants

Binding of antibody CSL362 hIgG1 to CD123 WT ECD and variants (analytes in solution) was performed at low (50nM) and high (300nM) concentration of analyte. Antibody CSL362 hIgG1 (captured ligand) was captured by Anti-Human Fc capture biosensor (AHC) (Sartorius, PN: 18-5060) for 300s at 0.5µg/mL. Analytes CD123 WT ECD and variants were titrated at 7 concentrations (1:2 dilution series) from 50nM to 0.78nM and from 300nM to 4.7nM. Association to analyte was monitored for 300s and dissociation for 600s or 900s. Double reference subtraction was performed against buffer only and biosensor loaded with a negative hIgG1 control. Regeneration was performed in 10mM Gly-HCl pH1.7. Data were analysed using the Octet Data Analysis software HT 12.0. Data were fitted to a 1:1 binding model. Kinetic rates ka and kd were globally fitted.

### 6H6 mIgG1 binding to CD123 WT ECD and variants

Binding of antibody 6H6 mIgG1 (Biolegend, PN: 306002; captured ligand) to CD123 WT ECD and variants (analytes in solution) was performed using Streptavidin capture biosensor (SA) (Sartorius, PN: 18-5019). CaptureSelect™ Biotin Anti-LC-kappa (Murine) (Thermo Fischer, PN: 7103152100) was captured for 600s at 1µg/mL on SA tips. Those biosensors were then used to capture antibody 6H6 mIgG1 for 300s at 2.5µg/mL. Analytes CD123 WT ECD and variants were titrated at 7 concentrations from 50 to 1.56nM. Association to analyte was monitored for 300s and dissociation for 600s. Buffer only well was used as reference. Regeneration was performed in 10mM Gly-HCl pH1.7. Data were analysed using the Octet Data Analysis software HT 12.0. Data were fitted (when possible) to a 1:1 binding model. Kinetic rates ka and kd were globally fitted.

### IL-3 binding to CD123 WT ECD and variants

Binding of IL-3 (Sino Biological, PN: 11858-H08H) (analyte in solution) to CD123 WT ECD and variants (captured ligands) was performed using Streptavidin capture biosensor (SA) (Sartorius, PN: 18-5019). CD123 WT ECD and variants were biotinylated using Biotinylation kit Type B (Abcam, PN: ab201796) following manufacturer instructions. Biotinylated CD123 WT ECD. and variants (ligands) were captured on SA tips for 1000s at 3µg/mL concentration. Analyte IL-3 was titrated at 7 concentrations from 500 to 7.8nM in PBS pH7.4. Association to analyte was monitored for 300s and dissociation for 120s. Buffer only well was used as reference. No regeneration was performed, and a new set of tips was used for each biotinylated captured ligand. Data were analysed using the Octet Data Analysis software HT12.0. Due to the fast on/off nature of the interaction data were analysed using Steady state analysis.

### Thermal stability analysis

Differential Scanning Fluorimetry (DSF) analysis were performed on a Bio-Rad CFX96 Touch Deep Well RT PCR Detection System. Sypro Orange 5000X in DMSO (Sigma, PN: S5692) was used at a final concentration of 5X. Temperature gradient was performed from 25 to 95°C in increment of 1.5°C in a reaction volume of 20μL. “FRET” scan mode was used to monitor fluorescence. All samples were analysed at a final concentration of 0.25mg/mL in triplicate in PBS pH7.4. The temperature of protein unfolding transition (Tm) was calculated using the 1st derivative method.

### Engineering of TF-1 cells expressing CD123 variants and functional assays thereof

RNPs were freshly prepared as outlined above. 50pmol ssDNA HDRT (180bp length, Ultramer DNA Oligonucleotides, synthetized by IDT) were added to the RNPs. Per reaction 0.2×10^6^ TF-1 cells were re-suspended in 10µl R buffer (Neon™ Transfection System) and electroporated using the Neon® Transfection System (1200V, 40ms, 1pulse). Following an expansion period of 12 days the edited cells were flow sorted based on the binding to MIRG123 and 6H6: WT (MIRG123^+^6H6^+^), KO (MIRG123^-^6H6^-^) and KI (MIRG123^-^6H6^+^).

To test responsiveness of TF-1 cells to human IL-3, 0.18×10^5^ sorted cells were distributed into white 96 well-plate clear bottom tissue-culture treated (Greiner) and different IL-3 concentrations indicated in **Fig. 5b** were added. In certain experiments MIRG123 was added in the concentrations shown in **Fig. 5c**. After 3 days, proliferation was assessed using CellTiter-Glo® 2.0 (Cat#G9241, Promega) according to the manufacturer’s protocol. Luminescence was assessed with the Synergy H1 (BioTek) with an integration time of 1s.

### Non-virally CRISPR/Cas9 mediated engineering of HSPCs

Leukopaks were purchased from CytoCare and HSPCs were isolated by the LP-34 Process using the CliniMACS Prodigy (Miltenyi). HSPCs were thawed in HSC-Brew GMP Basal Medium (Miltenyi) supplemented with HSC-Brew GMP Supplement, 2% human serum albumin, 100ng/mL Stem Cell Factor (SCF), 100ng/mL Thrombopoietin (TPO), 100ng/mL Fms-like tyrosine kinase 3 ligand (Flt3L) and 60ng/mL IL-3 (Miltenyi) at a concentration of 0.5×10^6^ cells/ml. Cells were electroporated 2 days later. gRNAs were freshly prepared as outlined above, but 50 µM crRNA and tracrRNA were used to form the gRNA and complexed with 1µM Spyfi Cas9 (Aldevron at 61.889 µM) at a molar ratio Cas9:gRNA = 1:2 and incubated for 20 min at RT. As control, incomplete RNPs lacking the site-specific crRNA were generated. During the RNPs complexing, HSPCs were collected, washed twice with electroporation buffer (Miltenyi) and resuspend in electroporation buffer at 1×10^6^ cells/90 µl. Cells were then mixed with 5µl RNP and ssDNA HDRT encoding the variants (5µl corresponding to 500pmol) and the whole volume was transferred into the electroporation nucleocuvette. Electroporation was performed with the CliniMACS Prodigy (600V 100μs burst / 400V 750μs square). Immediately after electroporation the cells were transferred to a 6 well plate and rested for 20 minutes at RT. After 20min, 2mL of pre-warmed HSC medium supplemented with 100ng/mL SCF, 100ng/mL TPO and 100ng/mL Flt3L was added and the plate was incubated at 37°C.

### CD123 and CCR5 gene editing with CRISPR/Cas9 and rAAV6 virus

#### CD34^+^ HSPC culture condition

Plerixafor mobilized peripheral blood CD34^+^ HSPCs were purchased from AllCells (Alameda, CA, USA). The cells were thawed per manufacturer’s instructions and were cultured at 37°C, 5% CO2, and 5% O2. Cell culture medium was GMP SCGM medium from CellGenix (Portsmouth, NH, USA) supplemented with human cytokine (PeproTech, Rocky Hill, NJ, USA) cocktail containing SCF 100ng/ml, TPO 100ng/ml, Flt3L 100ng/ml, and IL-6 100ng/ml. UM171 (35nM) (StemCell Technologies, Vancouver, Canada), streptomycin (20mg/ml) and penicillin (20U/ml) were added into the cell culture medium.

#### CD123 and CCR5 gene editing procedure

HSPCs were cultured and pre-stimulated for 72 hours after thaw and before gene editing. Chemically modified CD123 sgRNA and CCR5 sgRNA were synthesized by Synthego Corporation (Redwood City, CA, USA). SpyFi Cas9 was purchased from Aldevron, LLC (Fargo, ND, USA). CD123-E51K and CD123-E51T donor rAAV6 virus was purchased from SignaGen Laboratories (Frederick, MD, USA). CCR5-KO donor rAAV6 virus was produced in HEK-293T cells and was purified with AAVpro Purification Kit (TakaRa, San Jose, CA, USA). Electroporation of the RNP complex was performed using the Lonza 4D-Nucleofector (Lonza Group Ltd, Alpharetta, GA, USA) in P3 Primary Cell Solution with program DZ-100. Donor rAAV6 virus was immediately dispensed onto electroporated cells at multiplicity of infection (MOI) of 2.5×10^3^ vector genomes (vg) per cell for CD123 and 5.0×10^3^ vg per cell for CCR5 based on the titres determined by ddPCR. The cells were then divided into two halves at 2.5×10^5^ cells/ml. One half was plated in SCGM medium supplemented with cytokines and 10ng/ml IL-3 (PeproTech, Rocky Hill, NJ, USA) as +IL-3 treatment. The other half was plated in SCGM medium supplemented with cytokines only as -IL-3 treatment. After incubation for 24 hours, a medium change was performed to remove residual rAAV6 virus. The CD34^+^ HSPCs were cultured for up to 8 days for quantification of gene editing events, CFU assays and pSTAT5 staining and FACS analysis.

#### Methylcellulose colony formation unit (CFU) assay of non-virally edited HSPCs

CFU assay was started at 72h post gene editing. For each condition, 1.1mL of semi-solid methylcellulose medium (StemCell Technologies) containing 500 cells were plated in a well of a SmartDish (StemCell Technologies) in duplicates. The cells were incubated at 37 °C, 5% O2 and 5% CO2 for 14 days. The resulting progenitor colonies were counted and scored with STEMVision analysis (StemCell Technologies, Seattle, WA, USA) per manufacturer’s instruction. Colonies were picked and subjected to sequencing. Shortly, colonies were washed with PBS and then resuspended in 25 µl DNA-QuickExtract solution. PCRs were performed and sent for sanger sequencing.

#### Methylcellulose colony formation unit (CFU) assay of AAV-edited HSPCs

CFU assay was started at 48h post gene editing. For each condition, 1.1mL of semi-solid methylcellulose medium (StemCell Technologies, Seattle, WA, USA) containing 300 cells and with 10ng/ml IL-3 (+IL-3 treatment) or without IL-3 (-IL-3 treatment) were plated in a well of a SmartDish (StemCell Technologies, Seattle, WA, USA) in duplicates. The cells were incubated at 37 °C, 5% O2 and 5% CO2 for 14 days. The resulting progenitor colonies were counted and scored with STEMVision analysis (StemCell Technologies, Seattle, WA, USA) per manufacturer’s instruction.

#### pSTAT5 staining of non-virally edited HSPCs

On day 3 post gene editing, all cells were stimulated with 10ng/ml IL-3 for 1h at 37°C and then were subjected to pSTAT5 and CD123 staining. Briefly, after the 1h incubation, cells were fixed in 4% paraformaldehyde and then were permeabilized with ice-cold methanol. The permeabilized cells were stained with Alexa 647 mouse-anti-human Stat5 (pY694), BV650 mouse-anti-human CD123 clone 6H6 antibody and Alexa 488 MIRG123 in FACS buffer at RT for 1h. After staining, cells were washed with FACS buffer and then subjected for FACS analysis to Fortessa.

#### pSTAT5 staining of AAV-edited HSPCs

On day 8 post gene editing, all cells were switched to growth medium without IL-3 for IL-3 starvation. After 1-day IL-3 starvation, for +IL-3 treatment, 10ng/ml IL-3 was added back to the cells which were originally cultured in +IL-3 medium. For -IL-3 treatment, IL-3 was not added into the cells which were originally cultured in -IL-3 medium. All cell cultures were incubated for 1h and then were subjected to pSTAT5 staining. Briefly, after 1h incubation, cells were lysed and fixed in BD Phosflow Lyse/Fix Buffer and then were permeabilized with BD Phosflow Perm Buffer III (BD Biosciences, San Jose, CA, USA) following manufacturer’s instruction. The permeabilized cells were stained with Alexa 647 Mouse-anti-human Stat5 (pY694) antibody in PBS at RT for 1h. Alexa 647 Mouse isotype IgG (BD Pharmingen, San Jose, CA, USA) was used as a negative control. K-562 cells which were not subjected to IL-3 treatment nor IL-3 stimulation was stained with the same pSTAT5 antibody as a positive control. After staining, cells were washed with PBS and then were subjected to CytoFLEX Flow Cytometer (Beckman Coulter Life Sciences, Brea, CA) for FACS analysis.

#### Differentiation of non-virally edited HSPCs in vitro

Three days after electroporation cells were resuspended in StemPro media (Gibco) containing StemPro Nutrients, LDL 50ng/ml, P/S 1%, Glutamine 1%, Flt3 20ng/ml, TPO 50ng/ml, IL-6 50ng/ml, IL-3 10ng/ml, IL-2 10ng/ml, IL-7 20ng/ml, EPO 3ng/ml, GM-CSF 20ng/ml, SCF 100ng/ml, and 2000 cells/well were plated in a 96 round bottom well plate. In some wells, CSL-362-Tesirine at a concentration of 10ng/mL was added. After 14 days cells were collected, stained for CD33, GlyA/CD235a, CD14, CD15 and CD123 and acquired on a Fortessa.

#### Mice

All animal work was performed in accordance with the federal and cantonal laws of Switzerland. Protocols were approved by the Animal Research Commission of the Canton of Basel-Stadt, Switzerland. All mice were housed in a specific pathogen-free (SPF) condition in accordance with institutional guidelines and ethical regulations. NBSGW (stock# 026622) female mice were purchased from Jackson Laboratories. The HSPCs were edited as described above. Two days after electroporation cells were collected and frozen in CS10 (Stem Cell Technologies). Cells were thawed at the day of injection, washed and resuspended in PBS at 10×10^6^ live cells/ml. Recipient NBSGW female mice (3 weeks old) were injected into the tail vein. Chimerism was analysed after 6 and 10 weeks in the blood by flow cytometry. Mice were euthanized 16 weeks after humanization.

For secondary transplant, NSG-SGM3 (stock#013062) female mice were purchased from Jackson Laboratories. Mice were irradiated the day before BM transplant with 200cGy. Primary transplant mice were euthanized, bone marrow was isolated, and half of the bone marrow re-injected into the new host. Mice from secondary transplant were euthanized 8 weeks after transplant.

#### In vitro TCE-mediated killing assay of non-virally engineered HSPCs

For the TCE killing assays the HSPCs were edited as described above. Autologous human T cells were isolated from PBMCs as abovementioned and cultured overnight in supplemented media without stimulation. Two days after electroporation edited HSPCs were co-cultured in a 96 U-bottom plate together with human effector T cells at an effector to target ratio of 3:1 and the CD3/CSL362-TCE at 100ng/ml for 72 h at 37 °C. In a set of experiments MOLM-14 (labelled with CTV according to the manufacturer’s instruction) were added to the autologous T cells and edited HSPCs at an effector to target to tumour ratio of 3:0.5:0.5 together with the CD3/CSL362-TCE (100ng/ml). Cytotoxic activity (specific killing and elimination of non-edited HSCs) was analysed by flow-cytometry. Specific killing was calculated as follows: (1-No. alive target cells with TCE/No. alive target cells without TCE)*100.

#### Computational off-target prediction

The Cas-OFFinder algorithm ^44^ (release 2.4.1) was used to search the human reference genome hg38 for potential off-target sites in silico, using the following parameters: maximum number of mismatches = 3; maximum number of DNA bulges = 1; maximum number of RNA bulges = 1; PAM = NGG. The human genome reference hg38 (GRCh38 Genome Reference Consortium Human Reference 38), was downloaded from the Golden Path repository at https://hgdownload.cse.ucsc.edu/goldenPath/hg38/chromosomes/. All chromosomes were used in the alignments. To account for redundant off-target sites, unique target regions were selected by grouping overlapping sites using bedtools cluster (v2.30.0) and selecting a representative alignment. The same off-target analysis was performed for two additional gRNAs targeting the VEGFA^45^ and HBB^41,46^ genes which served as benchmarks.

#### Discover-Seq

Cells were nucleofected with an RNP complex to target either CD123 or BFP (used as a negative control). Thirteen hours after nucleofection, an aliquot of nucleofected cells were transferred to new media and genomic DNA was extracted 5 days later to check for genome editing efficiency. Ten million nucleofected cells were fixed in 1% formaldehyde at room temperature for 15 minutes. The fixation reaction was quenched with glycine to a final concentration of 125 mM. Cells were harvested and washed twice with chilled PBS and pellets were snap frozen and stored at -80° C until processing. To process, cell pellets were thawed on ice and incubated with lysis buffer 1 (LB) 1 (50 mM Hepes–KOH, pH 7.5; 140 mM NaCl; 1 mM EDTA; 10% Glycerol; 0.5% NP-40 or Igepal CA-630; 0.25% Triton X-100; 1X protease inhibitors) on ice for 10 minutes. Cells were pelleted by centrifugation and incubated in LB 2 (10 mM Tris–HCL, pH8.0; 200 mM NaCl; 1 mM EDTA; 0.5 mM EDTA; 1X protease inhibitors) for 5 minutes on ice. The extracted nuclei were pelleted by centrifugation and resuspended in LB 3 (10mM Tris–HCl, pH8; 100mM NaCl; 1mM EDTA; 0.1% NaDeoxycholate; 0.5% N-lauroylsarcosine; 1X protease inhibitors). Nuclei were sonicated using a Covaris S2 sonicator with the following settings: duty cycle 5%, intensity 5, 200 cycles per burst, 7 minutes. Debris were pelleted by centrifugation at 4°C and the supernatant was transferred to a 5mL tube. 100μL of Dynabeads protein A that had been prebound with MRE11 antibody (Novus NB 100-142) were added to the cell lysate and samples were incubated at 4°C overnight with rotation. Beads were collected on a magnetic stand and washed with ice-cold RIPA buffer 6 times, followed by a final wash with TBS before resuspending beads in 200 µL of elution buffer (50 mM Tris–HCl, pH 8; 10 mM EDTA; 1% SDS). The bead slurries were incubated overnight at 65°C to reverse crosslinks. Samples were treated with 1 mg/mL RNaseA (Ambion, catalog 2271) for 30 minutes at 37°C, followed by proteinase K treatment 20 mg/mL (Invitrogen, catalog 25530-049) for 1 hour at 55°C. DNA was then purified using a MinElute PCR Purification Kit (Qiagen, catalog #28004) and sequencing libraries were prepared using a NEBNext Ultra II kit (NEB, catalog E7645L). Samples were sequenced with 50-bp paired-end reads on an Illumina NextSeq at a depth of 30 million reads per sample. Following sequencing, data was analysed using the BLENDER2 pipeline (available on GitHub).

#### rhAmpSeq

Validation of off-target sites was performed using the RNase H-dependent amplification and sequencing (rhAmpSeq) system from IDT. rhAmpSeq primer panels for targeted amplification were generated using the rhAmpSeq design tool defining the insert size between 150-250bp. rhAmpSeq CRISPR library was prepared according to manufacturer’s instructions and sequenced on an Illumina MiniSeq instrument (MiniSeq Mid Output Kit, 300-cycles). Sequencing data were analysed with the rhAmpSeq CRISPR Analysis tool from IDT using the default settings.

#### CAST-Seq

Cells were nucleofected with RNP complexes to target CD123. Genomic DNA was extracted 5 days later using the NucleoSpin Tissue kit (Machery and Nagel). CAST-Seq library preparations were performed as described^42^, and data analysed using an improved bioinformatics pipeline^43^.

### Statistical Analysis

Statistical Analysis was performed on Prism 9.1.2 software (GraphPad). Number of donors and replicates are found within each figure legend.

