## Supplemental Figures and Tables for "Epitope Engineered Human Haematopoietic Stem Cells are Shielded from CD123-targeted Immunotherapy"

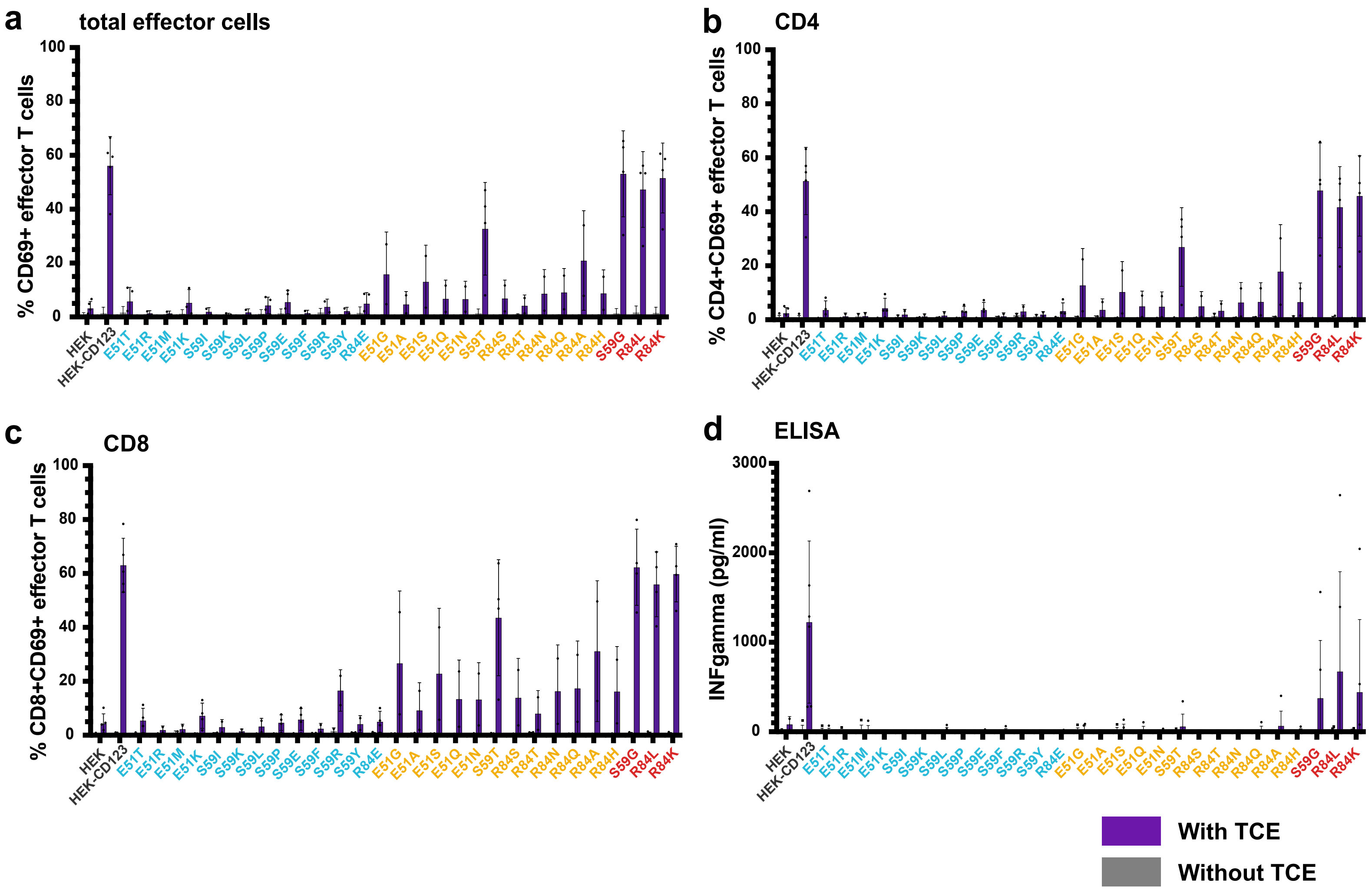


**Extended data Fig. 1**

72 hours co-culture of human effector T cells with HEK, HEK-CD123 and CD123 variants (E:T = 10:1) in the presence of the CSL362/OKT3-TCE (300ng/ml). Summary of flow cytometry CD69$^+$ total (**a**), gated CD4$^+$ (**b**), and CD8$^+$ (**c**) effector T cells after 3 days co-culture with (purple) and without (grey) TCE. Data from 5 independent donors and experiments with 2 technical replicates per group. **d,** IFNgamma secretion measured by ELISA in co-culture supernatants at 72h. **a-d,** Data from 4 blood donors and experiments with 2 technical replicates per group. Error bars: mean ± SD.

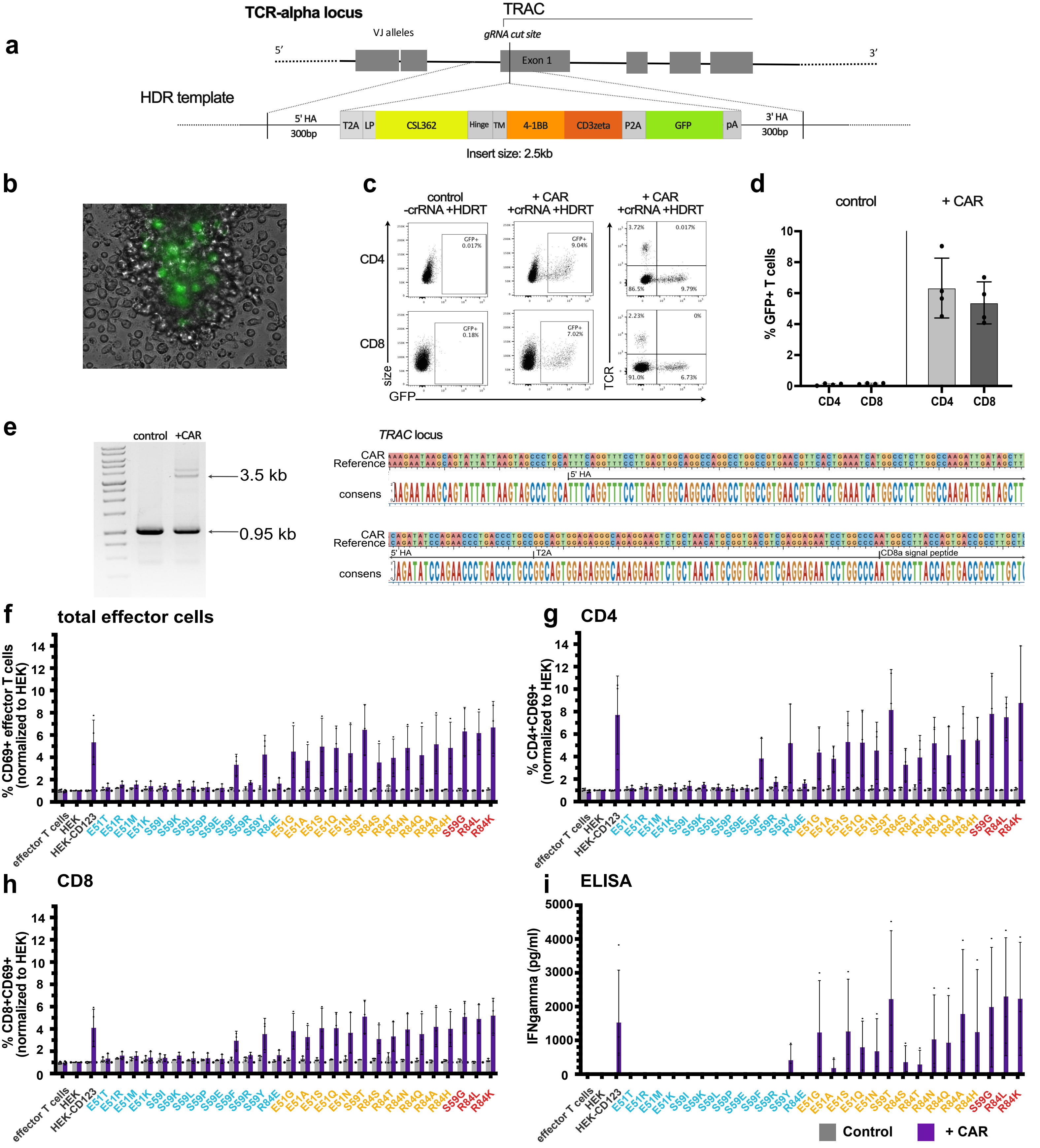


**Extended data Fig. 2**

**a,** Non-viral HDR-mediated integration of the CD123-specific second-generation CAR into Exon 1 of the *TRAC* locus using CRISPR/Cas9. **b,** Representative microscopy image (magnification 40x) day 4 post-electroporation showing GFP⁺ cells expressing the CAR-encoding template. **c,** Flow cytometry plots highlighting CAR insertion into the *TRAC* locus represented by fluorescent intensity of GFP with disrupted endogenous TCR expression in primary human gated CD4⁺ and CD8⁺ T cells. **d,** Mean knock-in efficiency of the CAR-encoding template in gated CD4⁺ and CD8⁺ T cells in “control” and “+CAR” cells at day 4-5. Data from 4 independent donors and experiments. **e, left** Gel image and **right** Sanger Sequencing results confirming correct HDRT integration at the *TRAC* locus in flow-sorted GFP⁺ CAR cells using PCR with primers annealing outside both arms of homology. Reference refers to the designed sequence in SeqBuilderPro (DNASTAR). **f-i,** 123CAR T cells (purple) or control cells (grey) were co-cultured with HEK, HEK-CD123 or its variants at an effector to target ratio of 10:1 for 24h. Summary of flow cytometry data indicating percentage of CD69⁺ total (**f**), gated CD4⁺ (**g**), or CD8⁺ (**h**) “control” (grey) or “+ CAR” (purple) T cells after 24h co-culture.**i.** Quantification of IFNg in supernatants of 24h co-cultures using ELISA. Error bars: mean ± SD. Data from 3 independent donors and experiments with 2 technical replicates per group.

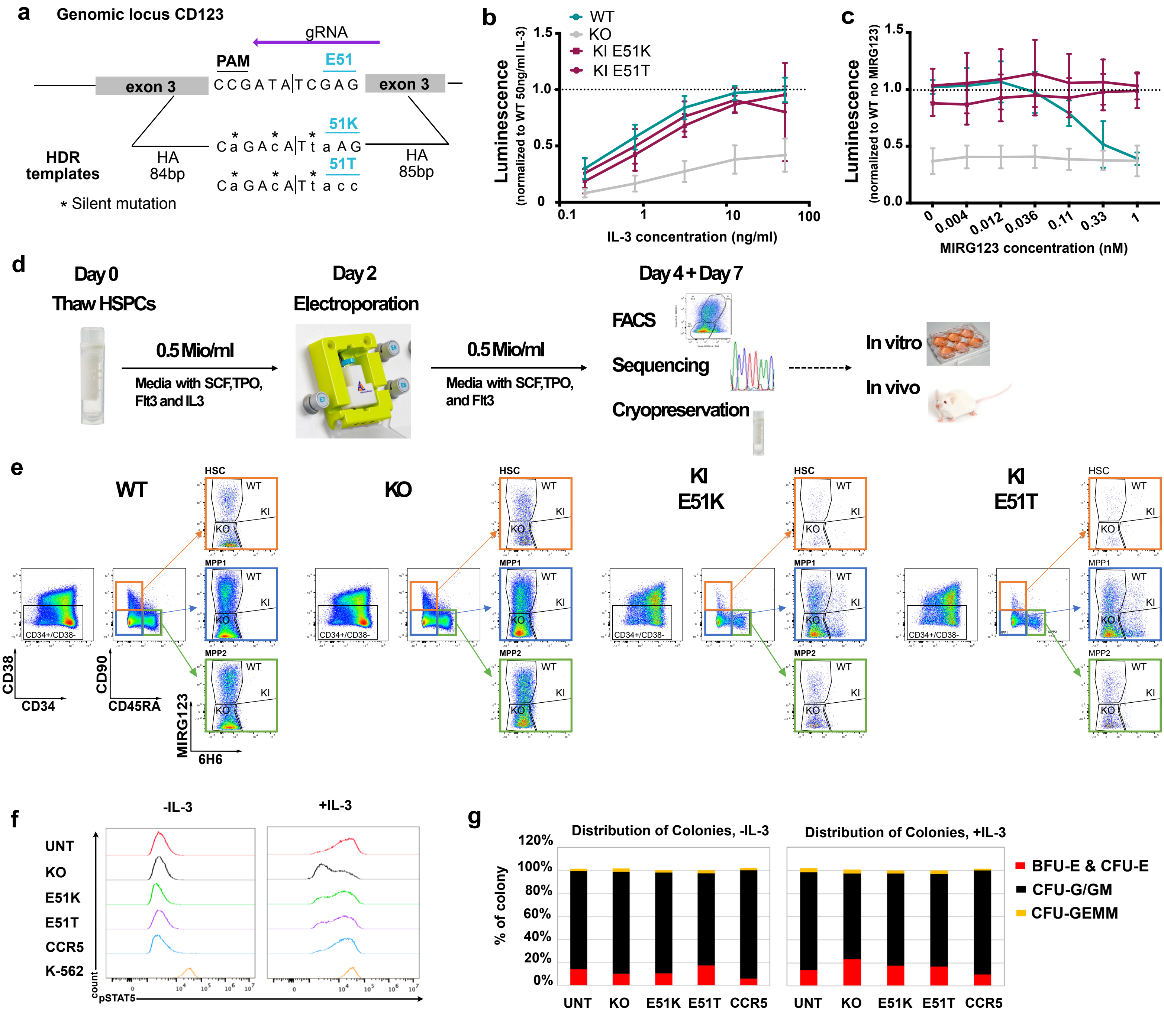


### Extended data Fig. 3

**a,** Schematic of HDR templates design for insertion of variants K and T at position 51. **b-c,** Engineered TF-1 cells were sorted based on the binding to the anti-CD123 mAbs 6H6 and MIRG123 (WT MIRG123$^+$6H6$^+$, KO MIRG123$^-$6H6$^-$, KI MIRG123$^-$6H6$^+$) and cultured for three days. **b,** with increasing concentrations of IL-3, or in **c,** with 2.5ng/ml IL-3 in the presence of increasing concentrations of MIRG123. Viable cells were quantified by luminescence and results normalized to WT cells cultured with 50ng/ml IL-3 (a) or to WT cells cultured without MIRG123 (b). Data from 4 independent experiments. **d,** Experimental design of non-viral CRISPR/Cas9-mediated HDR engineering of mobilized CD34$^+$ enriched peripheral blood HSPCs using GMP-compatible protocol with the CliniMACS Prodigy (Miltenyi). **e,** Gating strategy to monitor CD123 expression in HSC (orange, CD34$^+$CD38$^-$CD90$^+$CD45RA$^-$), multipotent progenitor 1 (MPP1; blue; CD34$^+$CD38$^-$CD90$^-$CD45RA$^-$) and MPP2 (green; CD34$^+$CD38$^-$CD90$^-$CD45RA$^+$) using the mAbs MIRG123 and 6H6 two and five days post electroporation. wt: MIRG123$^+$6H6$^+$, KO: MIRG123$^-$6H6$^-$, KI: MIRG123$^-$6H6$^+$ **f,** Representative histogram of phosphorylated STAT5 upon exposure to IL-3 in AAV6-edited HSPCs. CCR5 KO was used as negative, and K-562 cells as positive control. Data represents two independent experiments. **g,** In vitro differentiation of AAV6-edited HSPCs with and without IL-3. Colony Forming Units (erythroid: BFU-E & CFU-E, granulocytes/monocytes: CFU-G/GM and myeloid progenitors: CFU-GEMM) were scored based on morphological characteristics. Data from two independent experiments.

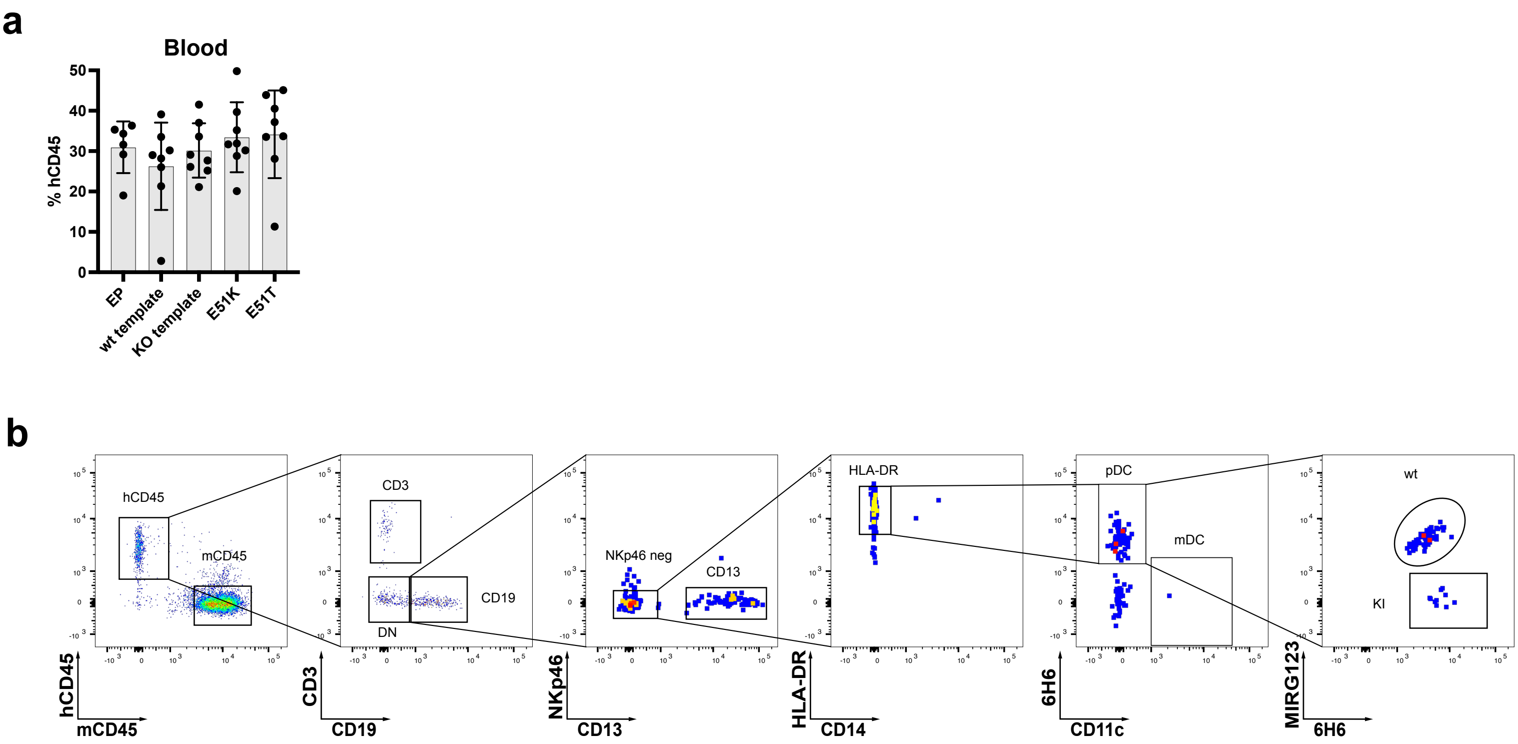


**Extended data Fig. 4**

**a,** Human chimerism (% hCD45⁺) in blood. **b,** Gating strategy to identify pDCs 16 weeks after injection of edited HSPCs in NBSGW mice.

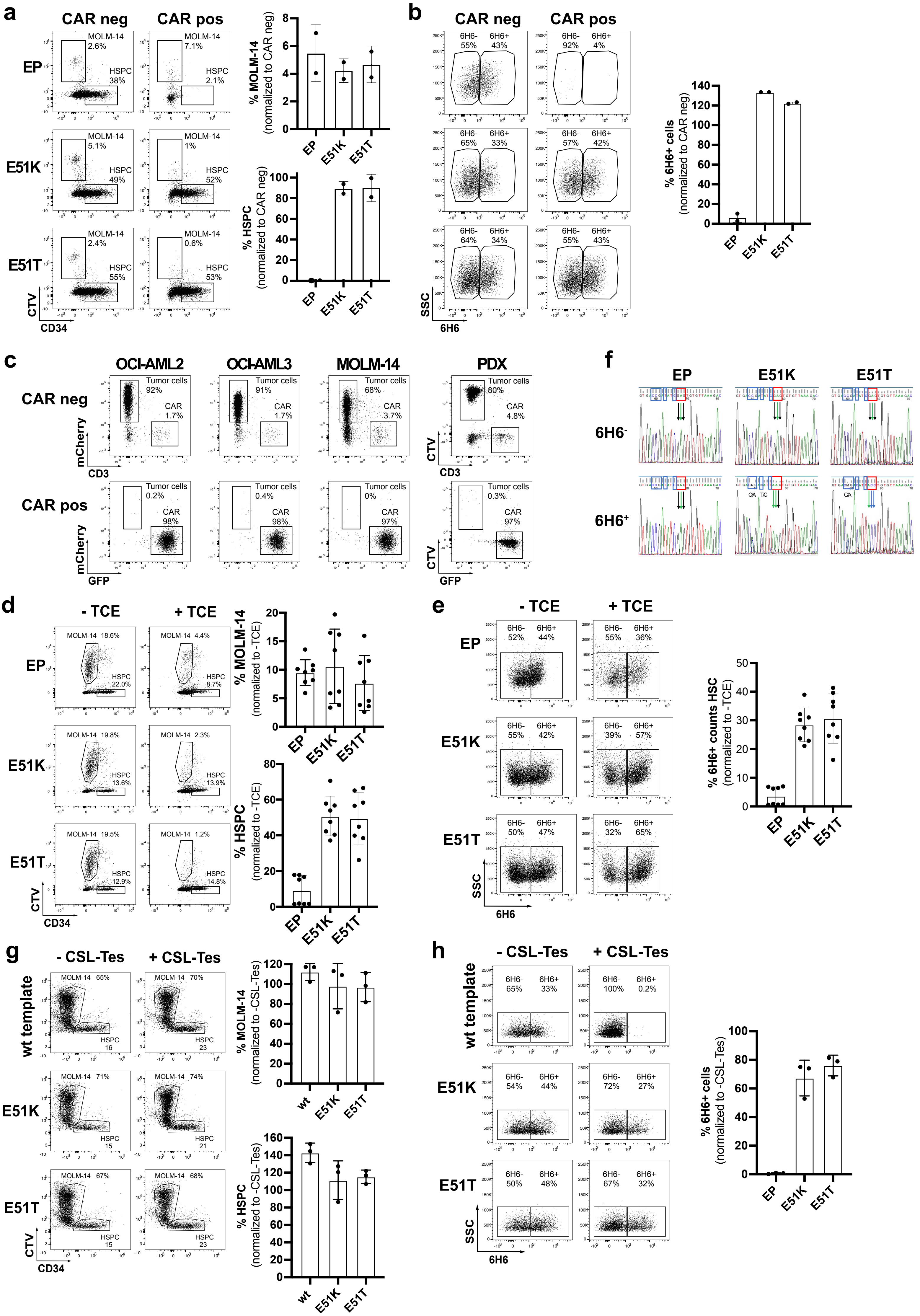

a
CAR neg
CAR pos
EP
E51K
E51T
MOLM-14
HSPC
CTV
CD34
% MOLM-14
(normalized to CAR neg)
% HSPC
(normalized to CAR neg)
b
6H6-
6H6+
SSC
6H6
% 6H6+ cells
(normalized to CAR neg)
c
OCI-AML2
OCI-AML3
MOLM-14
PDX
Tumor cells
CAR
mCherry
CD3
GFP
f
d
- TCE
+ TCE
% MOLM-14
(normalized to -TCE)
% HSPC
(normalized to -TCE)
e
% 6H6+ counts HSC
(normalized to -TCE)
g
- CSL-Tes
+ CSL-Tes
wt template
wt
% MOLM-14
(normalized to -CSL-Tes)
% HSPC
(normalized to -CSL-Tes)
h
% 6H6+ cells
(normalized to -CSL-Tes)

**Extended data Fig 5.**

**a-b,** Non-virally edited HSPCs co-cultured with MOLM-14 (CTV-labelled) and control T cells (CAR neg) or 123CAR for 3 days. **a, left** Representative dot plots indicating proportion (%) of MOLM-14 cells and non-virally edited $CD34^+$ HSPCs on day 3 of co-culture. **right** Quantification based on absolute counts. **b, left** FACS plots illustrating proportion (%) of wt or ki HSPCs at the end of the co-culture based on the binding characteristics to the mAb 6H6. Only clone 6H6 was used to avoid epitope masking by the 123CAR. **right** Quantification based on absolute counts. **c,** 2-day co-culture of AML cells MOLM-14-mCherry, OCI-AML2-mCherry, OCI-AML3-mCherry and PDX (CTV-labelled) with control T cells (CAR neg) or 123CAR. **d-e,** Non-virally edited HSPCs co-cultured with MOLM-14 (CTV-labelled) and autologous T cells with or without CSL362/OKT3-TCE (100ng/ml) for 3 days. Control condition (EP) are electroporated but non-edited HSPCs. Data from 2 individual donors performed in 3 independent experiments with 1 or 2 replicates per experiment. **d, left** Representative dot plot indicating proportion (%) of MOLM14 cells and $CD34^+$ HSPCs in different conditions on day 3 of co-culture. **right** Specific TCE-mediated killing of MOLM-14 or HSPCs at day 3. **e**, **left** FACS plots illustrating proportion (%) of edited HSPCs at the end of the co-culture based on the binding characteristics to the mAb 6H6. Only clone 6H6 was used to avoid epitope masking by the CSL362/OKT3-TCE. **right** Quantification based on absolute counts. **f,** Sanger Sequencing chromatogram of FACS-sorted $6H6^+$ and $6H6^-$ HSPCs on day 3 of co-culture with autologous T cells and CSL362/OKT3-TCE. Blue boxes: silent mutations, red boxes: E51K and E51T amino acids substitutions. **g-h,** Non-virally edited HSPCs co-cultured with MOLM-14 (CTV-labelled) with or without CSL362-Tesirine (10nM) for 3 days. Control condition (EP) are electroporated but non-edited HSPCs. Data from 2 individual donors performed in 3 independent experiments with 1 or 2 replicates per experiment. **g, left** Representative dot plot indicating proportion (%) of MOLM14 cells and $CD34^+$ HSPCs in different conditions on day 3 of co-culture. **right** Specific ADC-mediated killing of HSPCs at day 3. **h**, **left** FACS plots illustrating proportion (%) of edited HSPCs at the end of the co-culture based on the binding characteristics to the mAb 6H6. Only clone 6H6 was used to avoid epitope masking by the CSL362-Tesirine. **right** Quantification based on absolute counts.

**Extended data table 1: Computational off-target prediction**

| | chr | start | end | n_b | s_b | strand | guide | target | mismatches | bulges | bulge_type | gRNA | cluster | bulge_pos | insertion | deletion | CFD_score | MIT_score | CFD_final | max_CFD_sc | max_MIT_score |
|---|---|---|---|---|---|---|---|---|---|---|---|---|---|---|---|---|---|---|---|---|---|
| OnT | chrX | 1345396 | 1345416 | . | . | - | GTCTTTAACACACTCGATATNNN | GTCTTTAACACACTCGATATCGG | 0 | 0 | X | sg2440 | 27825 | NA | NA | NA | 1 | 1 | 1 | 1 | 1 |
| | chrY | 1345396 | 1345416 | . | . | - | GTCTTTAACACACTCGATATNNN | GTCTTTAACACACTCGATATCGG | 0 | 0 | X | sg2440 | 29440 | NA | NA | NA | 1 | 1 | 1 | 1 | 1 |
| OT1 | chr17 | 32136676 | 32136697 | . | . | - | GTCTTT-AACACACTCGATATNNN | GcCTTTCAACACACTCGtTATTGG | 2 | 1 | DNA | sg2440 | 10117 | 7 | NA | C | 0.11428571 | 0 | 0 | 0.08080808 | 0 |
| OT2 | chr10 | 61350729 | 61350748 | . | . | + | GTCTTTAACACACTCGATATNNN | GTtTTTAA-ACACTgGATATTGG | 2 | 1 | RNA | sg2440 | 2816 | 9 | C | NA | 0.034375 | 0.03209585 | 0.01427885 | 0.0175 | 0.03209585 |
| OT3 | chr7 | 30072218 | 30072237 | . | . | + | GTCTTTAACACACTCGATATNNN | G-CTTTAACACACTgGATAcTGG | 2 | 1 | RNA | sg2440 | 23825 | 2 | T | NA | 0.00454545 | 0 | 0.00433239 | 0.00433239 | 0 |
| OT4 | chr4 | 107801720 | 107801739 | . | . | + | GTCTTTAACACACTCGATATNNN | GTCTTTAACACACTCcAT-TCGG | 1 | 1 | RNA | sg2440 | 18980 | 19 | A | NA | 0 | 0 | 0 | 0 | 0 |
| OT5 | chr9 | 9095633 | 9095652 | . | . | + | GTCTTTAACACACTCGATATNNN | GTCTTTAAtACACTC-ATgTAGG | 2 | 1 | RNA | sg2440 | 26728 | 16 | G | NA | 0.328125 | 0 | 0 | 0 | 0 |
| OT6 | chr9 | 124360002 | 124360021 | . | . | + | GTCTTTAACACACTCGATATNNN | GTCTTTAACACACaC-cTATAGG | 2 | 1 | RNA | sg2440 | 27711 | 16 | G | NA | 0.1092437 | 0 | 0 | 0 | 0 |

**Extended data table 2: DISCOVER-Seq**

| Chr:Start-End | Cutsite | Discoscore | Cutsite Ends | Strand | PAM | Guide sequence | Mismatches | |
|---|---|---|---|---|---|---|---|---|
| chrX:1345397-1345416 | 1345399 | 207 | 192 | antisense | NGG | GTCTTTAACACACTCGATAT | 0 | **OnT** |
| chr4:98393545-98393564 | 98393547 | 9 | 4 | antisense | NGG | TTCTGTTACACTTTCAGTAT | 7 | OT7 |
| chr5:76852262-76852281 | 76852264 | 9 | 3 | antisense | NAG | TTTTTTATTACATTTGATAG | 7 | OT8 |
| chr6:117017325-117017344 | 117017327 | 9 | 9 | antisense | NAG | GTCCTTAACCCACTTTTTGA | 7 | OT9 |
| chr17:74873985-74874004 | 74873987 | 8 | 3 | antisense | NGG | GCCTTTAACTCCTTGGAACT | 7 | OT10 |
| chr11:77447109-77447128 | 77447125 | 7 | 3 | sense | NAG | TTCTATAACACATCCGCTAT | 5 | OT11 |
| chr1:230302954-230302973 | 230302956 | 7 | 3 | antisense | NAG | GGCTTTAACACACATTAATT | 6 | OT12 |
| chr2:209774760-209774779 | 209774762 | 7 | 3 | antisense | NGG | TGCTTTAAGAAACTAGATAA | 6 | OT13 |
| chr6:104499843-104499862 | 104499845 | 7 | 5 | antisense | NAG | GTCTCTAACATACTATTTTT | 6 | OT14 |
| chr11:123716422-123716441 | 123716424 | 7 | 7 | antisense | NAG | TTTTTTAACAAAATGGAAAG | 7 | OT15 |
| chr2:161232988-161233007 | 161232990 | 7 | 4 | antisense | NAG | TTTTTTAAAAAACTCATTTT | 7 | OT16 |
| chr5:164782381-164782400 | 164782397 | 7 | 4 | sense | NAG | TTCTTTTATTCAGTTTATAT | 7 | OT17 |
| chr14:23671996-23672015 | 23672012 | 7 | 6 | sense | NAG | CTCTTGAAGATACTTGATGG | 7 | |
| chr15:44003424-44003443 | 44003426 | 7 | 3 | antisense | NAG | GGTTTTAACAACCTACATCT | 7 | |
| chr1:160876572-160876591 | 160876588 | 6 | 5 | sense | NGG | GCCATTAACATTCTAGCTAC | 7 | |
| chr1:37513364-37513383 | 37513380 | 6 | 3 | sense | NAG | GTCCTAAAAACAAACGTTAC | 7 | |
| chr4:23452581-23452600 | 23452583 | 6 | 6 | antisense | NAG | GTATTTAGCAAAATAAAAAT | 7 | |
| chr4:38282453-38282472 | 38282469 | 6 | 6 | sense | NAG | ATTCTTAACACTCTCCTTTT | 7 | |
| chr4:46768054-46768073 | 46768056 | 6 | 6 | antisense | NGG | TGCTGTAGCACACGCCATTT | 7 | |
| chr5:75574555-75574574 | 75574557 | 6 | 5 | antisense | NAG | GTCTTAAACACATTTTACTA | 7 | |
| chr6:42392536-42392555 | 42392538 | 6 | 4 | antisense | NAG | GTCTTACACACACAGAATTT | 6 | |
| chr6:42811139-42811158 | 42811155 | 6 | 5 | sense | NAG | GTATTTTAGATATTAGATAT | 6 | |
| chr7:91072231-91072250 | 91072247 | 6 | 6 | sense | NGG | GGTTGTCAGACACCCTATAT | 7 | |
| chr9:68777861-68777880 | 68777863 | 6 | 3 | antisense | NAG | GTCTACAAAAAAATTAATAT | 7 | |
| chr9:86170488-86170507 | 86170504 | 6 | 3 | sense | NAG | TTCTTTAATACATTTTTTAA | 7 | |
| chr12:5414020-5414039 | 5414022 | 6 | 3 | antisense | NGG | ATATTAAACATAATTGATCT | 7 | |
| chr13:101500635-101500654 | 101500637 | 6 | 4 | antisense | NAG | GTTACTAACCAACCCAATAT | 7 | |
| chr15:35102144-35102163 | 35102160 | 6 | 4 | sense | NGG | GTCTTGAACCCAGGAGGTAA | 7 | |
| chr16:11111612-11111631 | 11111628 | 6 | 3 | sense | NAG | GTCCGTAAGAAAATCCATTT | 7 | |
| chrX:116397509-116397528 | 116397525 | 6 | 6 | sense | NAG | TACTTGTAGATTCTCGATAT | 7 | |
| chr1:150396163-150396182 | 150396179 | 5 | 4 | sense | NGG | GTCCTTAACTCACTTTTTGA | 7 | |
| chr1:198967994-198968013 | 198968010 | 5 | 3 | sense | NAG | GTTTTTACTACGTGGGATAT | 7 | |
| chr1:228239661-228239680 | 228239663 | 5 | 4 | antisense | NAG | GTCTTCAAAACGATCACTAA | 7 | |
| chr1:24872436-24872455 | 24872438 | 5 | 5 | antisense | NAG | GCCTTGGACACCCTCCTTTT | 7 | |
| chr1:73348984-73349003 | 73348986 | 5 | 5 | antisense | NAG | TCCATAATCACTCTGGATAT | 7 | |
| chr1:76267802-76267821 | 76267804 | 5 | 3 | antisense | NGG | TTCTGTAAGAAACCCTATCT | 7 | |
| chr2:125241448-125241467 | 125241464 | 5 | 3 | sense | NGG | GTTTTTAACCAACCAGGTAT | 6 | |
| chr2:151497647-151497666 | 151497663 | 5 | 5 | sense | NGG | GTGTTTGACTCTTTCCATCT | 7 | |
| chr2:158537676-158537695 | 158537692 | 5 | 4 | sense | NGG | ATGTCTGACAAACTTGATAC | 7 | |
| chr2:161902289-161902308 | 161902291 | 5 | 3 | antisense | NAG | GTTTTGAAAGCACTTTATAT | 6 | |
| chr2:21368644-21368663 | 21368646 | 5 | 4 | antisense | NGG | TCCTTTAACACAATCAAGCA | 7 | |
| chr2:213990843-213990862 | 213990845 | 5 | 5 | antisense | NGG | CTGTTTTACAAATTCCATAT | 6 | |
| chr2:217534119-217534138 | 217534121 | 5 | 4 | antisense | NAG | GTCAATGACAGACTGCATAT | 6 | |
| chr2:23727568-23727587 | 23727570 | 5 | 3 | antisense | NGG | GTTTTCAACTCATTGGGTAT | 6 | |
| chr2:37558013-37558032 | 37558015 | 5 | 4 | antisense | NAG | TTCACTCACATAGTTGATAT | 7 | |
| chr2:4207058-4207077 | 4207060 | 5 | 4 | antisense | NGG | GTAATTAACATACGCCAGAG | 7 | |
| chr2:46599864-46599883 | 46599880 | 5 | 3 | sense | NGG | CTCTTGCTCACACTTCACAT | 7 | |
| chr2:61633392-61633411 | 61633408 | 5 | 5 | sense | NGG | ATCTTTTACAATATACATAT | 7 | |
| chr2:83826419-83826438 | 83826435 | 5 | 3 | sense | NAG | TTCTTTTTCACCATCACTAT | 7 | |
| chr2:95516089-95516108 | 95516091 | 5 | 5 | antisense | NAG | ATCTTCCTCACACTCAATCT | 6 | |
| chr2:99592109-99592128 | 99592125 | 5 | 4 | sense | NGG | GGGTTAAAAACCCTCAAAAT | 7 | |
| chr3:131306513-131306532 | 131306515 | 5 | 3 | antisense | NAG | AGCTCTAGCCCACTCAACAT | 7 | |
| chr3:172065452-172065471 | 172065454 | 5 | 3 | antisense | NGG | CTATTTCACACACAGGAAGT | 7 | |
| chr4:110801722-110801741 | 110801724 | 5 | 3 | antisense | NAG | TCCTTTGAAAGACTGGATAA | 7 | |
| chr4:113827888-113827907 | 113827890 | 5 | 3 | antisense | NGG | TGATTTTGCCCACTCTATAT | 7 | |
| chr4:121189574-121189593 | 121189576 | 5 | 5 | antisense | NAG | GTAATTAAGACACCTGAACT | 7 | |
| chr4:122599865-122599884 | 122599867 | 5 | 3 | antisense | NGG | GTCCTTAGCAGTCAAGATAT | 6 | |
| chr4:128948874-128948893 | 128948876 | 5 | 4 | antisense | NAG | CTTTTCAAAATACTTGATAA | 7 | |

| | | | | | | | |
|---|---|---|---|---|---|---|---|
| chr4:160724372-160724391 | 160724374 | 5 | 4 | antisense | NAG | ATTTCTACCATATTCCATAT | 7 |
| chr4:39704462-39704481 | 39704464 | 5 | 4 | antisense | NAG | GTCTTTGTAACACTGTATAC | 6 |
| chr5:102466634-102466653 | 102466636 | 5 | 5 | antisense | NAG | CTCTTTTACACTCTTGGTGG | 7 |
| chr5:124181356-124181375 | 124181372 | 5 | 3 | sense | NAG | GTGTTTCCCACAGTCTATTC | 7 |
| chr5:144702366-144702385 | 144702368 | 5 | 4 | antisense | NAG | GCCTTTTCCAGTATCAATAT | 7 |
| chr5:166211956-166211975 | 166211972 | 5 | 3 | sense | NGG | GTCTTCCACACCCATGACTT | 7 |
| chr5:31519695-31519714 | 31519697 | 5 | 5 | antisense | NGG | GTCTTTAACATGATGAAGAA | 7 |
| chr5:36611659-36611678 | 36611675 | 5 | 4 | sense | NGG | GTCATGAAAGCATTAGATAA | 7 |
| chr5:54623367-54623386 | 54623369 | 5 | 5 | antisense | NAG | CTCTTGAAGACACTCGTGAA | 6 |
| chr5:85129060-85129079 | 85129076 | 5 | 4 | sense | NAG | GTCTATAAATTCAACGATAT | 7 |
| chr5:96395377-96395396 | 96395379 | 5 | 3 | antisense | NGG | TTGTTTATAATACTCCATAG | 7 |
| chr6:132125654-132125673 | 132125670 | 5 | 3 | sense | NAG | GTCTTTAAAACCATAACTTT | 7 |
| chr6:140907678-140907697 | 140907680 | 5 | 5 | antisense | NGG | TTCTTCTACTCAGTTGTTAT | 7 |
| chr6:156169137-156169156 | 156169139 | 5 | 4 | antisense | NGG | GTCTGTGTTACTCTTGGTAT | 7 |
| chr6:77607872-77607891 | 77607874 | 5 | 3 | antisense | NGG | GTCTTTGTCAGACTAAATTA | 7 |
| chr7:153983450-153983469 | 153983466 | 5 | 5 | sense | NAG | AACTTTAAGCCAGTAGATCT | 7 |
| chr7:91572035-91572054 | 91572037 | 5 | 4 | antisense | NGG | GCCTTCAACATACTAGCATT | 7 |
| chr7:9283010-9283029 | 9283012 | 5 | 4 | antisense | NAG | CTCTTTGGGCCACTCTCTAT | 7 |
| chr8:112756372-112756391 | 112756374 | 5 | 4 | antisense | NGG | CTCTTTAAAATACTCCTATT | 7 |
| chr8:129413389-129413408 | 129413405 | 5 | 3 | sense | NGG | GTCTAGTACTCACCAGACAT | 7 |
| chr8:138032367-138032386 | 138032369 | 5 | 4 | antisense | NGG | GTATTTTAAATTCTAAATAT | 7 |
| chr8:14012870-14012889 | 14012886 | 5 | 4 | sense | NAG | GTATTTAGCACATACTTTCT | 7 |
| chr8:41302503-41302522 | 41302519 | 5 | 5 | sense | NAG | TTCTGTAAGAAATCCGATCT | 7 |
| chr8:58334163-58334182 | 58334179 | 5 | 3 | sense | NAG | GACTTTAATACCCTCACAGT | 7 |
| chr8:61603716-61603735 | 61603732 | 5 | 4 | sense | NGG | GTACTTCACACATTAGAAAG | 7 |
| chr9:76727131-76727150 | 76727147 | 5 | 4 | sense | NAG | GGCCTTGACCCACTAGATGC | 7 |
| chr10:116445064-116445083 | 116445066 | 5 | 3 | antisense | NAG | GATTTTACTACACTGCATTT | 7 |
| chr10:128992782-128992801 | 128992784 | 5 | 3 | antisense | NAG | TTCTTTGAGAAAGTGGATAA | 7 |
| chr10:22171999-22172018 | 22172015 | 5 | 5 | sense | NAG | GTATTTATTCCACTTCATCT | 7 |
| chr11:44589383-44589402 | 44589399 | 5 | 3 | sense | NGG | GTTTTTAACACAGGAAAGCT | 7 |
| chr12:51774641-51774660 | 51774657 | 5 | 4 | sense | NAG | GTCTCAAAGGCCCTTGTTAT | 7 |
| chr12:73253333-73253352 | 73253335 | 5 | 3 | antisense | NGG | GTCTTCTTCACAGGCCAAAT | 7 |
| chr12:77016358-77016377 | 77016360 | 5 | 5 | antisense | NAG | ATCTTTTACACTTTTGATTG | 7 |
| chr12:79862546-79862565 | 79862548 | 5 | 4 | antisense | NGG | GTTTTTAGCTTTCTTGCTAT | 7 |
| chr12:91160175-91160194 | 91160191 | 5 | 4 | sense | NAG | GACATTAACAAACAAAAAAT | 7 |
| chr14:31448219-31448238 | 31448235 | 5 | 3 | sense | NAG | AGCTTCAGCACACTTGCTAT | 6 |
| chr15:50545240-50545259 | 50545256 | 5 | 3 | sense | NAG | GTTAATAAAACAATAGCTAT | 7 |
| chr16:11381527-11381546 | 11381543 | 5 | 5 | sense | NAG | GTCTTTCAGACAGCAAAAAT | 7 |
| chr16:17289676-17289695 | 17289678 | 5 | 3 | antisense | NAG | GTTTTTAATATTCTGAATAG | 7 |
| chr16:80625037-80625056 | 80625053 | 5 | 3 | sense | NAG | GAATTTAACAGAATTGATGA | 7 |
| chr18:26660389-26660408 | 26660405 | 5 | 3 | sense | NAG | ATCTTTAAGAAATTCGGAAA | 7 |
| chr18:49219005-49219024 | 49219007 | 5 | 3 | antisense | NGG | GTCTTTATTAGTCTTGCTAG | 7 |
| chr18:64636662-64636681 | 64636678 | 5 | 5 | sense | NAG | CTCTTTACCAGACACCGTAT | 6 |
| chr19:23023176-23023195 | 23023192 | 5 | 5 | sense | NGG | TTATAGAACACACTAAAGAT | 7 |
| chr1:115063246-115063265 | 115063248 | 4 | 3 | antisense | NGG | GTCTTTTGCCCACTTTTTAA | 7 |
| chr1:13122490-13122509 | 13122506 | 4 | 4 | sense | NAG | GTTTTCAACATACAACATTT | 7 |
| chr1:173723621-173723640 | 173723637 | 4 | 3 | sense | NAG | GTCCTTTGCAGGCTAGATAT | 6 |
| chr1:175263656-175263675 | 175263658 | 4 | 3 | antisense | NGG | GTCTTTAATTTACCCTAAGT | 7 |
| chr1:194676897-194676916 | 194676913 | 4 | 3 | sense | NAG | TTCTTTAAATATCTACATAT | 7 |
| chr1:240888808-240888827 | 240888810 | 4 | 3 | antisense | NGG | GTGGTTAACTGAGTCGATTC | 7 |
| chr1:246213452-246213471 | 246213468 | 4 | 4 | sense | NAG | ATCTTTGAGAAAATGGATAG | 7 |
| chr1:247400413-247400432 | 247400429 | 4 | 3 | sense | NAG | GTGTTTCTCAAAAGCGACAT | 7 |
| chr1:31949842-31949861 | 31949858 | 4 | 3 | sense | NAG | ATCATTAATCCAGTGGATAG | 7 |
| chr1:56748124-56748143 | 56748140 | 4 | 3 | sense | NAG | TTCTTGTAGATTCTGGATAT | 7 |
| chr1:93877062-93877081 | 93877078 | 4 | 3 | sense | NGG | TTCTTTAAGAGATTCAACAT | 6 |
| chr1:9568001-9568020 | 9568003 | 4 | 4 | antisense | NAG | CTCTTCAATACAGTAGCCAT | 7 |
| chr2:110654566-110654585 | 110654568 | 4 | 4 | antisense | NGG | ATCCTTCAAAAAGTCAATAT | 7 |
| chr2:12908049-12908068 | 12908051 | 4 | 4 | antisense | NGG | CTCAACAACACAGGCTATAT | 7 |
| chr2:143108472-143108491 | 143108474 | 4 | 3 | antisense | NAG | ATCTTTAGAAAATTTAATAT | 7 |
| chr2:15975361-15975380 | 15975377 | 4 | 3 | sense | NGG | GTCGAAAACATTCTAGAAAT | 7 |
| chr2:220371861-220371880 | 220371877 | 4 | 4 | sense | NGG | GTGTGTATGTCACTTGATTT | 7 |

| | | | | | | | |
|---|---|---|---|---|---|---|---|
| chr2:230857634-230857653 | 230857636 | 4 | 4 | antisense | NAG | ATCCTTAACACTATGGTTGT | 7 |
| chr2:31908354-31908373 | 31908356 | 4 | 3 | antisense | NGG | TTTTTTTACTCTCACCATAT | 7 |
| chr2:37056190-37056209 | 37056206 | 4 | 4 | sense | NAG | GTCCACAATACAACAGATAT | 7 |
| chr2:43430785-43430804 | 43430801 | 4 | 4 | sense | NAG | GGCTTCTACCCACTAGATGC | 7 |
| chr2:58511603-58511622 | 58511605 | 4 | 3 | antisense | NAG | TTTTTTCAATCACTCCAGAT | 7 |
| chr2:76140-76159 | 76142 | 4 | 3 | antisense | NGG | TACTTTATCCCATTAGAAAT | 7 |
| chr2:88209548-88209567 | 88209564 | 4 | 3 | sense | NGG | GGCTTAAAAACCTTCCATAC | 7 |
| chr3:138192938-138192957 | 138192954 | 4 | 3 | sense | NGG | GTGTTTTAAAAGCTCTTTAT | 7 |
| chr3:191179702-191179721 | 191179704 | 4 | 3 | antisense | NGG | GAAATAAACCCACTCCAAAT | 7 |
| chr3:35209265-35209284 | 35209267 | 4 | 4 | antisense | NAG | GTCTTTTACCCTTAGAATAT | 7 |
| chr3:43207624-43207643 | 43207640 | 4 | 4 | sense | NAG | GGCTTTAACACAGAAGGGCT | 7 |
| chr3:70685392-70685411 | 70685408 | 4 | 4 | sense | NAG | GTTCTTCATAGACTTGATGT | 7 |
| chr4:124542687-124542706 | 124542703 | 4 | 4 | sense | NAG | TTGTTTAAGACAGTCAATCA | 7 |
| chr4:126925949-126925968 | 126925951 | 4 | 4 | antisense | NAG | TTCTTTTAAATTCTGGATAT | 6 |
| chr4:129576610-129576629 | 129576626 | 4 | 4 | sense | NAG | GTATGAAACCCACTTGATCA | 7 |
| chr4:150459549-150459568 | 150459565 | 4 | 4 | sense | NGG | GACAATAACACATTCCTTAC | 7 |
| chr4:163419507-163419526 | 163419523 | 4 | 3 | sense | NGG | TTCTTTAAATTACTAAATTT | 7 |
| chr4:48513776-48513795 | 48513792 | 4 | 3 | sense | NAG | GATTTTAACACATATACTAT | 7 |
| chr4:785181-785200 | 785197 | 4 | 3 | sense | NGG | GTTTTTAAAAGACACGACCC | 7 |
| chr4:86399425-86399444 | 86399441 | 4 | 4 | sense | NAG | GTCTTTTGCCCACTTTTTAG | 7 |
| chr5:123550178-123550197 | 123550180 | 4 | 4 | antisense | NGG | TGCTTGAAACCACTATATAT | 7 |
| chr5:142723610-142723629 | 142723626 | 4 | 4 | sense | NGG | GTCTTTCTCCTACTCCATGG | 7 |
| chr5:162484621-162484640 | 162484623 | 4 | 3 | antisense | NAG | GTCTAAAACATAGTGAATAA | 7 |
| chr5:170868972-170868991 | 170868988 | 4 | 3 | sense | NAG | GTTTTTAGTATACCAGGTAT | 7 |
| chr5:29099568-29099587 | 29099584 | 4 | 4 | sense | NGG | GACTTGCACACAGTCACTTT | 7 |
| chr5:60625855-60625874 | 60625857 | 4 | 3 | antisense | NAG | AGCTTGAATACACACAATAG | 7 |
| chr5:64584446-64584465 | 64584448 | 4 | 4 | antisense | NAG | GACTTTTCCATACTGGGTGT | 7 |
| chr5:85149665-85149684 | 85149681 | 4 | 3 | sense | NAG | GTTTTTAAAATAATTGTTGT | 7 |
| chr6:126369597-126369616 | 126369613 | 4 | 4 | sense | NGG | GTCCTTAGCCCACTTTTTGT | 7 |
| chr6:156004686-156004705 | 156004702 | 4 | 4 | sense | NAG | GAGATTGTCCCACTCCATAT | 7 |
| chr6:22808448-22808467 | 22808464 | 4 | 3 | sense | NGG | AACTAAATCACACTGGATGT | 7 |
| chr6:37631603-37631622 | 37631619 | 4 | 4 | sense | NGG | GACATCCACACCCTTGATTT | 7 |
| chr6:51169251-51169270 | 51169267 | 4 | 3 | sense | NGG | ATATTGAAAACAACAGATAT | 7 |
| chr6:85680235-85680254 | 85680237 | 4 | 4 | antisense | NGG | TTTTTTAATACAATAGAGAC | 7 |
| chr6:96345167-96345186 | 96345183 | 4 | 4 | sense | NAG | CTTTTTAAGATTCTACATAT | 7 |
| chr7:100092525-100092544 | 100092541 | 4 | 3 | sense | NAG | GACTTGCACACAGGCAGTAT | 7 |
| chr7:101254781-101254800 | 101254797 | 4 | 4 | sense | NAG | GTATCTAAGACACCCCATAT | 5 |
| chr7:129888973-129888992 | 129888975 | 4 | 4 | antisense | NGG | GTCCAAATCACACTCTGTTT | 7 |
| chr7:133807293-133807312 | 133807295 | 4 | 3 | antisense | NAG | GACTTTCACAGACACTCTAT | 6 |
| chr7:16098659-16098678 | 16098675 | 4 | 4 | sense | NAG | ACCTTCAACAGACTTGTTAT | 6 |
| chr7:68485664-68485683 | 68485666 | 4 | 3 | antisense | NAG | GTCATGGACTCACTTGCCAT | 7 |
| chr7:85137458-85137477 | 85137474 | 4 | 3 | sense | NAG | CTCTTTATCAAAGTTGTCAT | 7 |
| chr7:94123559-94123578 | 94123575 | 4 | 4 | sense | NGG | GTCTGGAAGAGACTGGATTC | 7 |
| chr8:29117752-29117771 | 29117768 | 4 | 3 | sense | NAG | TTTTCAAACATACTCAAAAT | 7 |
| chr8:29784115-29784134 | 29784131 | 4 | 3 | sense | NAG | CTTCTTATAACACTTTATAT | 7 |
| chr8:43341146-43341165 | 43341148 | 4 | 4 | antisense | NAG | GTTTTTAAGAGATATGATAG | 7 |
| chr8:52421166-52421185 | 52421182 | 4 | 3 | sense | NAG | GAATTTAACAAATTCAAAAC | 7 |
| chr8:61066765-61066784 | 61066781 | 4 | 3 | sense | NAG | GTCAATGACACACAACATTT | 7 |
| chr8:62530966-62530985 | 62530982 | 4 | 4 | sense | NAG | TGCTTCAACTCACTCCCTAT | 6 |
| chr8:65228932-65228951 | 65228934 | 4 | 3 | antisense | NGG | TTGTTTAACTAACTACCTAT | 7 |
| chr9:110894300-110894319 | 110894316 | 4 | 4 | sense | NGG | TTCTTTTCCTTACTGCATAT | 7 |
| chr9:116822270-116822289 | 116822286 | 4 | 4 | sense | NGG | TTCTTCAAGACATTTGAGAC | 7 |
| chr9:137546316-137546335 | 137546332 | 4 | 3 | sense | NAG | GTCTTGTATCCAATAAATAT | 7 |
| chr9:1681589-1681608 | 1681605 | 4 | 3 | sense | NAG | GACTACAACACAATAGCTGT | 7 |
| chr9:80241420-80241439 | 80241436 | 4 | 4 | sense | NGG | GTCTTTAATAAACTTTATTG | 6 |
| chr9:95080887-95080906 | 95080889 | 4 | 3 | antisense | NAG | GTCCTTAAGAATCTCTGTAA | 7 |
| chr10:112032964-112032983 | 112032980 | 4 | 3 | sense | NAG | TTCTTTGGCCCACTTGAGGT | 7 |
| chr10:130923936-130923955 | 130923952 | 4 | 3 | sense | NGG | ATCTCTCTCAGACTTGATTT | 7 |
| chr10:34783155-34783174 | 34783171 | 4 | 3 | sense | NGG | GAGTATAACACACTGAATGA | 7 |
| chr10:37241468-37241487 | 37241484 | 4 | 4 | sense | NGG | CTCTTTAACAAACTTTTTAG | 6 |
| chr10:57040614-57040633 | 57040616 | 4 | 3 | antisense | NAG | GTATTTTAGATACCTCATAT | 7 |

| | | | | | | | |
|---|---|---|---|---|---|---|---|
| chr10:83349358-83349377 | 83349360 | 4 | 3 | antisense | NAG | CCTTTTTAGACACTAGACAT | 7 |
| chr11:104172054-104172073 | 104172056 | 4 | 4 | antisense | NGG | TTTTTTAACACATTTTATAT | 5 |
| chr11:126783443-126783462 | 126783459 | 4 | 3 | sense | NAG | GACTTCAACATATTATATTT | 7 |
| chr11:16721109-16721128 | 16721111 | 4 | 4 | antisense | NAG | GCCTTTAACTTTCTCTGTCT | 7 |
| chr11:25531491-25531510 | 25531493 | 4 | 3 | antisense | NAG | GACTTCCACAAAGTGGATAA | 7 |
| chr11:80722576-80722595 | 80722592 | 4 | 4 | sense | NAG | TTCTGTAACACAGACTGAAT | 7 |
| chr11:98284379-98284398 | 98284395 | 4 | 4 | sense | NAG | AGCTTTAAATAACCCGTTAT | 7 |
| chr12:117225488-117225507 | 117225504 | 4 | 4 | sense | NAG | GGCTTCCACCCACTAGATGC | 7 |
| chr12:1872113-1872132 | 1872129 | 4 | 3 | sense | NGG | GAGTTTAACATCCTAGATGT | 6 |
| chr13:26187921-26187940 | 26187937 | 4 | 3 | sense | NAG | GTGTAAAGCACACTGGACAC | 7 |
| chr13:56073933-56073952 | 56073949 | 4 | 4 | sense | NGG | GATTTTACCATATTCAATAT | 6 |
| chr13:78982231-78982250 | 78982247 | 4 | 3 | sense | NGG | TTGTTTCTAACACTCAAAAT | 7 |
| chr14:38900537-38900556 | 38900539 | 4 | 4 | antisense | NAG | GTCTTTGACATAATTTTAAT | 7 |
| chr15:54379779-54379798 | 54379795 | 4 | 3 | sense | NAG | TTCTTTTACATTCTCTGTAT | 6 |
| chr15:97339594-97339613 | 97339596 | 4 | 4 | antisense | NGG | AGCTTTAACGTACACATTAT | 7 |
| chr16:6548322-6548341 | 6548338 | 4 | 3 | sense | NAG | GTGCTTAAAAGAATAGAAAT | 7 |
| chr16:79712986-79713005 | 79713002 | 4 | 4 | sense | NAG | GACTTTCAGACCCCTGATCT | 7 |
| chr17:42273843-42273862 | 42273859 | 4 | 3 | sense | NAG | GATTTGAAAAAACTCGACTT | 7 |
| chr18:35753244-35753263 | 35753260 | 4 | 4 | sense | NAG | GACTTTATTCAACTCATTAT | 7 |
| chr18:3984994-3985013 | 3984996 | 4 | 4 | antisense | NGG | GGCTTTAACAAAAACAATGA | 7 |
| chr18:54691383-54691402 | 54691399 | 4 | 4 | sense | NGG | GTCTACAACTAAATCCCTAT | 7 |
| chr18:74642385-74642404 | 74642401 | 4 | 3 | sense | NGG | GTTTTTACCATATGAGTTAT | 7 |
| chr18:79934954-79934973 | 79934956 | 4 | 3 | antisense | NGG | GTTTATAAAACCCTCAAATT | 7 |
| chr19:23037580-23037599 | 23037582 | 4 | 3 | antisense | NAG | TTGTTCAACTCATTGCATAT | 7 |
| chr19:44381504-44381523 | 44381506 | 4 | 3 | antisense | NAG | GTTCTGAAAACATTCTATAT | 6 |
| chr20:39024050-39024069 | 39024052 | 4 | 3 | antisense | NAG | GTCTACTGCACAGTGGATGT | 7 |
| chr20:39641180-39641199 | 39641182 | 4 | 3 | antisense | NAG | GTCTGGAACATACTCATTAT | 5 |
| chr20:9881754-9881773 | 9881770 | 4 | 4 | sense | NAG | TCCTTTTATGCTCTGGATAT | 7 |
| chr21:22438530-22438549 | 22438532 | 4 | 4 | antisense | NAG | ATCTCTTAAATTCTCAATAT | 7 |
| chrX:11761989-11762008 | 11762005 | 4 | 4 | sense | NAG | GCCTTTAACATACTTTTAAT | 6 |
| chr9:98682327-98682346 | 98682343 | 3 | 3 | sense | NGG | ATATTTACCACACACGAGAT | 5 |
| chr14:99905851-99905870 | 99905867 | 3 | 3 | sense | NAG | GCCTTTAACCCACTCTGTCT | 5 |

**Extended data table 3: CAST-Seq**

| Target | Function | | ID | Sequence (5'-3') |
|---|---|---|---|---|
| CD123 | CAST-Seq PCRI | bait | 8278 | TTTTAGATCCAAACCCACCAATC |
| | CAST-Seq PCRI | decoy fwd | 8217 | CCGGTAAATCATACTCTCTATTGTT |
| | CAST-Seq PCRI | decoy rev | 8216 | CATAGAATAGTCGGCGTCTT |
| | CAST-Seq PCRII | bait nested | 8215 | GACTGGAGTTCAGACGTGTGCTCTTCCGATCTCCAAACCCACCAATCACGAACC |
| Linker primers | CAST-Seq PCRI | prey | 4032 | GTAATACGACTCACTATAGGGC |
| | CAST-Seq PCRII | prey nested | 4033 | ACACTCTACACTCTTTCCCTACACGACGCTCTTCCGATCTAGGGCTCCGCTTAAGGGAC |
| Linker oligos | positive strand | | 4038 | GTAATACGACTCACTATAGGGCTCCGCTTAAGGGACT |
| | negative strand | | 4039 | P-GTCCCTTAAGCGGAGC-NH3 |

**Extended data Table 4 - Sequences / Flow Cytometry Antibodies**

**Sequences**

| NAME | | SEQUENCE (5'->3') |
|---|---|---|
| **crRNA** | | |
| crRNA TRAC | | AGAGTCTCTCAGCTGGTACA |
| crRNA CD123 | | GTCTTTAACACACTCGATATCGG |
| crRNA CCR5 | | GCAGCATAGTGAGCCCAGAAGGG |
| **HDRT** | | |

| | | |
|---|---|---|
| HDR template 123CAR | | TTTCAGGTTTCCTTGAGTGGCAGGCCAGGCCTGGCCGTGAACGTTCACTGAAATCATGGCCTCTTGGCCAAGATTGATAGCTTGTGCCTGTCCCTGAGTCCCAGTCCATCACGAGCAGCTGGTTTCTAAGATGCTATTTCCCGTATAAAGCATGAGACCGTGACTTGCCAGCCCCACAGAGCCCCGCCCTTGTCCATCACTGGCATCTGGACTCCAGCCTGGGTTGGGGCAAAGAGGGAAATGAGATCATGTCCTAACCCTGATCCTCTTGTCCCACAGATATCCAGAACCCTGACCCTGCCGGCAGTGGAGAGGGCAGAGGAAGTCTGCTAACATGCGGTGACGTCGAGGAGAATCCTGGCCCAATGGCCTTACCAGTGACCGCCTTGCTCCTGCCGCTGGCCTTGCTGCTCCACGCCGCCAGGCCGgacatcgtgatgacacagtctccagattccctggccgtgagcctgggagagagggctaccatcaattgcgagtccagccagagcctgctgaactctggcaatcagaagaactatctgacatggtaccagcagaagcctggccagcccccta agccactgatctattgggcctccaccagagagtctggagtgccagaccggttctctggatccggcagcggcacagacttcaccctgacaatctcttccctgcaggccgaggacgtggccgtgtactattgccagaacgattactcttatccctacaccttcggccagggcacaaagctggagatcaagggtggcggagggtctggcggtgggggatccggaggcggtgggagcgaggtgcagctggtgcagtctggagctgaggtgaagaagccaggcgagagcctgaagatctcttgcaagggcagcggctactcttttacagactactatatgaagtgggctaggcagatgcctggcaagggcctggagtggatgggcgatatcatcccaagcaacggcgccaccttctacaatcagaagtttaagggccaggtgacaatctctgctgacaagagcatctctaccacatatctgcagtggtccagcctgaaggccagcgataccgctatgtactattgtgctaggtcccacctgctgagagcttcctggttcgcttactggggccagggcaccatggtgacagtgtcttccACCACGACGCCAGCGCCGCGACCACCAACACCGGCGCCCACCATCGCGTCGCAGCCCCTGTCCCTGCGCCCAGAGGCGTGCCGGCCAGCGGCGGGGGGCGCAGTGCACACGAGGGGGCTGGACTTCGCCTGTGATATCTACATCTGGGCGCCCTTGGCCGGGACTTGTGGGGTCCTTCTCCTGTCACTGGTTATCACCCTTTACTGCAAACGGGGCAGAAAGAAACTCCTGTATATATTCAAACAACCATTTATGAGACCAGTACAAACTACTCAAGAGGAAGATGGCTGTAGCTGCCGATTTCCAGAAGAAGAAGAAGGAGGATGTGAACTGAGAGTGAAGTTCAGCAGGAGCGCAGACGCCCCCGCGTACCAGCAGGGCCAGAACCAGCTCTATAACGAGCTCAATCTAGGACGAAGAGAGGAGTACGATGTTTTGGACAAGAGACGTGGCCGGGACCCTGAGATGGGGGGAAAGCCGCAGAGAAGGAAGAACCCTCAGGAAGGCCTGTACAATGAACTGCAGAAAGATAAGATGGCGGAGGCCTACAGTGAGATTGGGATGAAAGGCGAGCGCCGGAGGGGCAAGGGGCACGATGGCCTTTACCAGGGTCTCAGTACAGCCACCAAGGACACCTACGACGCCCTTCACATGCAGGCCCTGCCCCCTCGCggcagcggcgccacaaacttctctctgctaaagcaagcaggtgatgttgaagaaaaccccgggcctGTGAGCAAGGGCGAGGAGCTGTTCACCGGGGTGGTGCCCATCCTGGTCGAGCTGGACGGCGACGTAAACGGCCACAAGTTCAGCGTGTCCGGCGAGGGCGAGGGCGATGCCACCTACGGCAAGCTGACCCTGAAGTTCATCTGCACCACCGGCAAGCTGCCCGTGCCCTGGCCCACCCTCGTGACCACCCTGACCTACGGCGTGCAGTGCTTCAGCCGCTACCCCGACCACATGAAGCAGCACGACTTCTTCAAGTCCGCCATGCCCGAAGGCTACGTCCAGGAGCGCACCATCTTCTTCAAGGACGACGGCAACTACAAGACCC |
| HDR template E51K | | TTTTAGATCCAAACCCACCAATCACGAACCTAAGGATGAAAGCAAAGGCTCAGCAGTTGACCTGGGACCTTAACAGAAATGTGACaGAcATtaagTGTGTTAAAGACGCCGACTATTCTATGCCGGTAAATCATACTCTCTATTGTTTTTTATTTTTATTTTATTTATTTATGTATTTA |
| HDR template E51T | | TTTTAGATCCAAACCCACCAATCACGAACCTAAGGATGAAAGCAAAGGCTCAGCAGTTGACCTGGGACCTTAACAGAAATGTGACaGAcATtaccTGTGTTAAAGACGCCGACTATTCTATGCCGGTAAATCATACTCTCTATTGTTTTTTTATTTTTATTTTATTTATTTATGTATTTA |

| **Primer** | | |
| --- | --- | --- |
| Fwd_HDRT CD123-CAR | | TTTCAGGTTTCCTTGAGTGGCA |
| Rev_HDRT CD123-CAR | | TGGCCATTCCTGAAGCAAGGA |
| Fwd_Sequencing 123CAR | | CTCCCATTCTGCTAATGCCCA |
| Rev_Sequencing 123CAR | | CTCCTGCCACCTTCTCTTCATC |
| hu CD123-Reverse primer | | TGAAGCTCATAGCGAAATTTGC |
| hu CD123-E51K-FWD | | TGTGACCGATATCAAGTGTGTTAAAGACGC |
| hu CD123-E51K-REV | | GCGTCTTTAACACACTTGATATCGGTCACA |
| hu CD123-E51T-Fwd | | TGTGACCGATATCTATTGTGTTAAAGACGC |
| hu CD123-E51T-Rev | | GCGTCTTTAACACAATAGATATCGGTCA |
| hu CD123-E51S-Fwd | | TGTGACCGATATCTCGTGTGTTAAAGACGC |
| hu CD123-E51S-Rev | | GCGTCTTTAACACACGAGATATCGGTCACA |
| hu CD123-E51Q-Fwd | | TGTGACCGATATCCAGTGTGTTAAAGACGC |
| hu CD123-E51Q-Rev | | GCGTCTTTAACACACTGGATATCGGTCACA |
| hu CD123-E51R-Fwd | | TGTGACCGATATCCGGTGTGTTAAAGACGC |
| hu CD123-E51R-rev | | GCGTCTTTAACACACCGGATATCGGTCACA |
| hu CD123-E51M-Fwd | | TGTGACCGATATCATGTGTGTTAAAGACGC |
| hu CD123-E51M-Rev | | GCGTCTTTAACACACATGATATCGGTCACA |
| hu CD123-E51G-Fwd | | TGTGACCGATATCGGGTGTGTTAAAGACGC |
| hu CD123-E51G-Rev | | GCGTCTTTAACACACCCGATATCGGTCACA |
| hu CD123-E51N-Fwd | | TGTGACCGATATCAACTGTGTTAAAGACGC |
| hu CD123-E51N-Rev | | GCGTCTTTAACACAGTTGATATCGGTCACA |
| hu CD123-E51A-Fwd | | TGTGACCGATATCGCGTGTGTTAAAGACGC |
| hu CD123-E51A-Rev | | GCGTCTTTAACACACGCGATATCGGTCACA |
| HuCD123-S59I-Fwd | | AGACGCCGACTATATTATGCCGGCAGTGAAC |
| HuCD123-S59I-Rev | | GTTCACTGCCGGCATAATATAGTCGGCGTCT |
| HuCD123-S59G-Fwd | | AGACGCCGACTATGGTATGCCGGCAGTGAAC |
| HuCD123-S59G-Rev | | GTTCACTGCCGGCATACCATAGTCGGCGTCT |
| HuCD123-S59P-Fwd | | AGACGCCGACTATCCTATGCCGGCAGTGAAC |
| HuCD123-S59P-Rev | | GTTCACTGCCGGCATAGGATAGTCGGCGTCT |
| HuCD123-S59E-Fwd | | AGACGCCGACTATGAGATGCCGGCAGTGAAC |

| | | |
|---|---|---|
| HuCD123-S59E-Rev | | GTTCACTGCCGGCATCTCATAGTCGGCGTCT |
| HuCD123-S59L-Fwd | | AGACGCCGACTATTTAATGCCGGCAGTGAAC |
| HuCD123-S59L-Rev | | GTTCACTGCCGGCATTAAATAGTCGGCGTCT |
| HuCD123-S59T-Fwd | | AGACGCCGACTATTATATGCCGGCAGTGAAC |
| HuCD123-S59T-Rev | | GTTCACTGCCGGCATATAATAGTCGGCGTCT |
| HuCD123-S59F-Fwd | | AGACGCCGACTATTTTATGCCGGCAGTGAAC |
| HuCD123-S59F-Rev | | GTTCACTGCCGGCATAAAATAGTCGGCGTCT |
| HuCD123-S59R-Fwd | | AGACGCCGACTATAGAATGCCGGCAGTGAAC |
| HuCD123-S59R-Rev | | GTTCACTGCCGGCATTCTATAGTCGGCGTCT |
| HuCD123-S59K-Fwd | | AGACGCCGACTATAAAATGCCGGCAGTGAAC |
| HuCD123-S59K-Rev | | GTTCACTGCCGGCATTTTATAGTCGGCGTCT |
| HuCD123-R84T-Fwd | | ACCAACTACACCGTCACAGTGGCCAACCCA |
| HuCD123-R84T-Rev | | TGGGTTGGCCACTGTGACGGTGTAGTTGGT |
| HuCD123-R84K-Fwd | | ACCAACTACACCGTCAAAGTGGCCAACCCA |
| HuCD123-R84K-Rev | | TGGGTTGGCCACTTTGACGGTGTAGTTGGT |
| HuCD123-R84S-Fwd | | ACCAACTACACCGTCTCAGTGGCCAACCCA |
| HuCD123-R84S-Rev | | TGGGTTGGCCACTGAGACGGTGTAGTTGGT |
| HuCD123-R84Q-fwd | | ACCAACTACACCGTCCAAGTGGCCAACCCA |
| HuCD123-R84Q-Rev | | TGGGTTGGCCACTTGGACGGTGTAGTTGGT |
| HuCD123-R84N-Fwd | | ACCAACTACACCGTCAATGTGGCCAACCCA |
| HuCD123-R84N-Rev | | TGGGTTGGCCACATTGACGGTGTAGTTGGT |
| HuCD123-R84E-Fwd | | ACCAACTACACCGTCGAAGTGGCCAACCC |
| HuCD123-R84E-Rev | | TGGGTTGGCCACTTCGACGGTGTAGTTGGT |
| HuCD123-R84H-Fwd | | ACCAACTACACCGTCCATGTGGCCAACCCA |
| HuCD123-R84H-Rev | | TGGGTTGGCCACATGGACGGTGTAGTTGGT |
| HuCD123-R84A-Fwd | | ACCAACTACACCGTCGCAGTGGCCAACCCA |
| HuCD123-R84A-Rev | | TGGGTTGGCCACTGCGACGGTGTAGTTGGT |
| HuCD123-R84L-Fwd | | ACCAACTACACCGTCCTAGTGGCCAACCCA |
| HuCD123-R84L-Rev | | TGGGTTGGCCACTAGGACGGTGTAGTTGGT |
| T7 promoter-huCD123 forward primer | | 1741<br>TAATACGACTCACTATAGG |

| | | |
|---|---|---|
| sequencing_gDNA_CD123_f | | TTCGAACTCCAACCTGTCACC |
| sequencing_gDNA_CD123_r | | CGGAAACGTCTTGTTCGTCA |

**Flow Cytometry Antibodies**

| *TARGET* | *FLUOROCHROME* | *CLONE* | *CONCENTRATI(* | *COMPANY* | *CAT. NR.* |
|---|---|---|---|---|---|
| CD123 | Biotin | CSL362 | 1:50 | - | - |
| CD123 | Alexa Fluor 650 | 6H6 | 1:200 | BioLegend | 306020 |
| Streptavidin | PE | | 1:500 | BioLegend | 405204 |
| Streptavidin | FITC | - | 1:200 | BioLegend | 405202 |
| CD3 | PerCPCy5.5 | UCHT1 | 1:100 | BioLegend | 300430 |
| CD3 | Brilliant Violet 421 | UCHT1 | 1:50 | BioLegend | 300434 |
| CD4 | APC | RPA-T4 | 1:100 | BioLegend | 300514 |
| CD4 | Alexa Fluor 488 | RPA-T4 | 1:100 | BioLegend | 300519 |
| CD4 | Alexa Fluor 510 | RPA-T4 | 1:100 | BioLegend | 300545 |
| CD8 | PE | RPA-T8 | 1:100 | BioLegend | 301008 |
| CD8 | Alexa Fluor 700 | RPA-T8 | 1:100 | BioLegend | 301028 |
| TCRa/b | Brilliant Violet 421 | IP26 | 1:25 | BioLegend | 306722 |
| CD33 | PE CF594 | WM53 | 1:100 | BD Biosciences | 562492 |
| CD69 | Alexa Fluor 605 | FN50 | 1:25 | BioLegend | 310938 |
| GlyA/CD235a | PE | 11E4B-7-6 | 1:100 | Beckman Coulter | A07792 |
| CD34 | PerCPCy5.5 | 8G12 (HPCA2) | 1:10 | BD Biosciences | 347222 |
| CD38 | Alexa Fluor 700 | LS198-4-3 | 1:50 | Beckman Coulter | B23489 |
| CD45RA | APC/Fire | HI100 | 1:50 | Biolegend | 304152 |
| CD90 | PE/Cyanine7 | 5E10 | 1:50 | BD Biosciences | 561558 |
| CD11c | PE CF594 | B-Ly6 | 1:50 | BD Biosciences | 562393 |
| CD14 | Brilliant Violet 711 | M5E2 | 1:50 | Biolegend | 301838 |
| CD14 | PE | M5E2 | 1:50 | Biolegend | 982508 |
| CD15 | APC/Cyanine7 | W6D3 | 1:50 | Biolegend | 323048 |
| NKp46/CD335 | APC | 9E2/NKp46 | 1:50 | BD Biosciences | 558051 |
| CD117 | Brilliant Violet 711 | 104D2 | 1:50 | Biolegend | 313230 |
| CD33 | FITC | HIM3-4 | 1:50 | Biolegend | 303304 |
| CD33 | Brilliant Violet 785 | WM35 | 1:50 | Biolegend | 303428 |
| CD19 | Brilliant Violet 785 | SJ25C1 | 1:50 | Biolegend | 363028 |
| CD13 | PE/Cyanine7 | WM15 | 1:50 | Biolegend | 301712 |
| HLA-DR | BUV496 | G46-6 | 1:50 | BD Biosciences | 749866 |
| FceRI | Alexa Fluor 488 | | 1:50 | Biolegend | 334640 |
| CD45 | V500 | HI30 | 1:200 | BD Biosciences | 560777 |
| CD45 | Brillant Violet 605 | 30-F11 | 1:200 | Biolegend | 103155 |
| CD45 | Alexa Fluor 700 | 30-F11 | 1:200 | Biolegend | 103128 |
| MIRG123-bio | | | | home made | |
| Zombie UV Fixable Viability Kit | - | | 1:1000 | BioLegend | 423108 |

| | | | | | |
|---|---|---|---|---|---|
| eBioscience Fixabile Viability Dye eFluor 780 | - | | 1:1000 | Invitrogen | 65-0865-14 |
| CellTrace Violet Cell Proliferation Kit | - | | 1:1000 | Invitrogen | C34557 |
| pSTAT5 | Alexa Fluor 647 | 47/Stat5(pY694) | 1:50 | BD Biosciences | 612599 |
| IgG1 isotype κ control | Alexa Fluor 647 | | | BD Biosciences | 557732 |
| FcR blocking reagent human | | | 1:200 | Miltenyi | 130-059-901 |
| CD16/32 | | 2.4G2 | 1:200 | BioXcell | |
